# Target-specific precision of CRISPR-mediated genome editing

**DOI:** 10.1101/387027

**Authors:** Anob M. Chakrabarti, Tristan Henser-Brownhill, Josep Monserrat, Anna R. Poetsch, Nicholas M. Luscombe, Paola Scaffidi

## Abstract

The CRISPR-Cas9 system has successfully been adapted to edit the genome of various organisms. However, our ability to predict editing accuracy, efficacy and outcome at specific sites is limited by an incomplete understanding of how the bacterial system interacts with eukaryotic genomes and DNA repair machineries. Here, we performed the largest comparison of indel profiles to date, examining over one thousand sites in the genome of human cells, and uncovered general principles guiding CRISPR-mediated DNA editing. We find that precision of DNA editing varies considerably among sites, with some targets showing one highly-preferred indel and others displaying a wide range of infrequent indels. Editing precision correlates with editing efficiency, homology-associated end-joining for both insertions and deletions, and a preference for single-nucleotide insertions. Precise targets and the identity of their preferred indel can be predicted based on simple rules that mainly depend on the fourth nucleotide upstream of the PAM sequence. Regardless of precision, site-specific indel profiles are highly robust and depend on both DNA sequence and chromatin features. Our findings have important implications for clinical applications of CRISPR technology and reveal general patterns of broken end-joining that can inform us on DNA repair mechanisms in human cells.

## INTRODUCTION

The CRISPR-Cas9 system has quickly become the preferred tool for genome engineering, enabling site-specific alterations in a variety of organisms and cellular contexts (Hsu et al., 2014). The system relies on the combined use of the bacterial Cas9 endonuclease and a single-guide RNA (sgRNA) either to substitute, insert or delete DNA sequences in almost any desired location in the genome (Hsu et al., 2014). Regardless of the experimental setting and application, genome editing by the CRISPR-Cas9 system entails three steps: i) scanning of the genome by the RNA-guided Cas9 nuclease (RGN) to find the DNA sequence complementary to the sgRNA, ii) creation of a DNA double strand break (DSB) by Cas9, and iii) repair of the lesion by the endogenous DNA repair machinery (Hsu et al., 2014). Both the accuracy and efficiency of the processes involved in each of these steps strongly affect the outcome of CRISPR-mediated editing and consequently the utility of the technology. Since the adaptation of the CRISPR system as an engineering tool, several studies have provided insights into the mechanisms affecting CRISPR-mediated DNA editing and have improved the technology (Brinkman et al., 2018; Henser-Brownhill et al., 2017; Horlbeck et al., 2016; Hsu et al., 2014; Isaac et al., 2016; van Overbeek et al., 2016; Tsai et al., 2015; Uusi-Mäkelä et al., 2018). However, fundamental questions about how the mammalian genome and proteins interact with Cas9 and the sgRNAs and how cells respond to CRISPR-induced DNA damage remain unanswered. Increasing our knowledge of the mechanisms regulating these interactions is crucial to maximise the potential and safety of CRISPR-based approaches.

A key prerequisite for a good editing tool is the ability to discriminate between on-target and homologous off-target sites. Characterisation of selected sgRNAs using both *in vitro* and cellular assays has provided important information about parameters influencing RGN specificity identifying the seed region of guide RNAs (the 10-to 12-nt sequence adjacent to the PAM sequence) as critical for recognition of target sequences (Hsu et al., 2014). This characterization has guided sgRNA-designing algorithms and improved CRISPR fidelity. However, systematic investigation of off-target cleavage sites has shown that predicting the specificity profile of any given RGN is not straightforward, and has revealed that our understanding of how RGNs scan the mammalian genome is incomplete (Tsai et al., 2015). Importantly, by showing that truncated guide RNAs (17-18nt) exhibit substantially reduced off-target DSBs, this large-scale analysis has proposed alterations that can considerably improve the technology and benefit various applications (Tsai et al., 2015). This example illustrates how systematic characterisation of CRISPR-induced alterations in experimental systems may inform us on how RGNs interact with complex genomes and help optimizing editing outcome.

In addition to specificity, another feature that can vary widely across RGNs is activity. While direct measurement of cleavage activity at a given target is not simple, sgRNA efficacy has been inferred either by quantifying the frequency of insertion/deletion (indel) formation or by evaluating the ability of an sgRNA to induce an expected phenotype. Analysis of large-scale studies has revealed sequence patterns correlating with sgRNA activity and has guided refinement of algorithms for sgRNA design (Doench et al., 2016; Wang et al., 2014). Although *in silico* predictions of sgRNA efficacy have improved considerably, concordance between predicted and empirically measured indel activity remains moderate (Henser-Brownhill et al., 2017). Thus, while we have achieved a qualitative understanding of RGN activity determinants, additional parameters not included in the current algorithms likely contribute to the overall outcome. The epigenetic status of target sequences may be one of such factors. Although correlative evidence and *in vitro* studies have implicated chromatin in the modulation of RGN activity (Horlbeck et al., 2016; Uusi-Mäkelä et al., 2018), formal demonstration that the chromatin status of an endogenous locus affects its editing potential is still lacking.

DSBs induced by RGNs at target sites are recognized by the cell’s DNA damage response pathways and repaired. When accurate repair fails, this process creates a chance to modify the endogenous sequence. When an exogenous repair template is provided, the homologous recombination repair pathway (HR) allows introduction of precise modifications in the DNA sequence, including single point mutations or insertion of exogenous sequences (Hsu et al., 2014). In the absence of a template, RGN-induced DSBs are often repaired through relatively error-prone mechanisms that result in insertions or deletions of variable length. Indels disrupting gene open reading frames lead to production of truncated, often non-functional proteins, making RGN-induced editing an effective means to induce gene knock-out (KO) (Hsu et al., 2014). Despite the wide use of the CRISPR system to generate KO alleles, our understanding of the mechanisms driving indel formation is still limited, making the functional outcome of genome editing unpredictable and often preventing a rational use of the technology. Based on the type of indels observed upon RGN-mediated editing, two major repair pathways have been implicated in the formation of RGN-induced indels: canonical non-homologous end joining (cNHEJ), known to induce small indels, and microhomology mediated end-joining (MMEJ), typically generating larger deletions at regions of microhomology (MH) (Deriano and Roth, 2013). Of note, genetic studies examining the general role of these pathways in the formation of CRISPR-mediated indels are currently lacking and the predominant method of repair of RGN-induced DSBs remains unclear. Based on the assumption that NHEJ is the main pathway involved in CRISPR-mediated indel formation, repair outcome was thought to be random. However, recent characterisation of indel patterns at multiple genomic locations revealed that individual targets show reproducible repair outcome, with distinct preferences for class (insertion or deletions) and size of indels (van Overbeek et al., 2016). This finding suggests a deterministic nature of RGN-induced break repair and raises questions about the factors involved in defining these non-random patterns. Here, we performed a large-scale genomic characterisation of indel patterns examining over one thousand sites in the genome of human cells, with the aim of understanding how genetic and epigenetic factors influence CRISPR-mediated DNA editing. We find that target sites display highly variable editing precision, i.e. the preference for a specific indel. This site-specific feature depends on both DNA sequence and chromatin states of the target sites and can be predicted. Our findings suggest that examination of a few target sites is insufficient to derive general principles of DNA editing, and that selection of a predictable target is an effective strategy to induce desired CRISPR-mediated alterations.

## RESULTS

### Large-scale analysis of indel patterns

To characterize general patterns of RGN-induced indels, we selected 1492 target sites across the genome and retrieved the corresponding sgRNAs from an arrayed lentiviral library that we had previously generated (Table S1) (Henser-Brownhill et al., 2017). The library targets 450 nuclear genes with multiple sgRNAs and has shown overall high activity (Henser-Brownhill et al., 2017). At least three sites for each gene were selected, taking care to space the target regions along genes (Fig. S1A) and using sgRNAs with high predicted activity (Fig. S1B). Retrieved sgRNAs were combined and sequenced to confirm homogeneous representation in the resulting pools (see Methods) (Fig. 1A and Fig. S1C). Pooled sgRNAs were then transduced into HepG2 cells expressing Cas9 and allowed to edit their target sites for 5 days, a timeframe found to be sufficient to generate detectable indels, but short enough to avoid KO-induced phenotypic changes that may confound the results (Fig. 1A, and Fig. S1 D,E). Upon isolation of genomic DNA, target regions were captured by pull-down using custom probes, and sequenced at ~6000 to 8000-fold coverage (Fig. 1A, Fig. S2 A,B). As expected, infection with pooled sgRNAs resulted in a high proportion of cells with unedited sequence at each target site, since only a small fraction of cells within the population expressed each sgRNA and could edit the corresponding site (Fig. S2A). Therefore, we developed a custom computational pipeline to filter background signal from unedited cells for a given sgRNA, which enabled robust detection of indels (see Methods) (Fig. S2A). 1122 sites showed detectable indels, ranging from 1 to 188 per target, with a median count of 32 (Fig. S2C). To avoid confounding results with low-count sites, only targets showing at least 10 indels (646 sites) were selected for downstream analysis.

**Figure 1.**
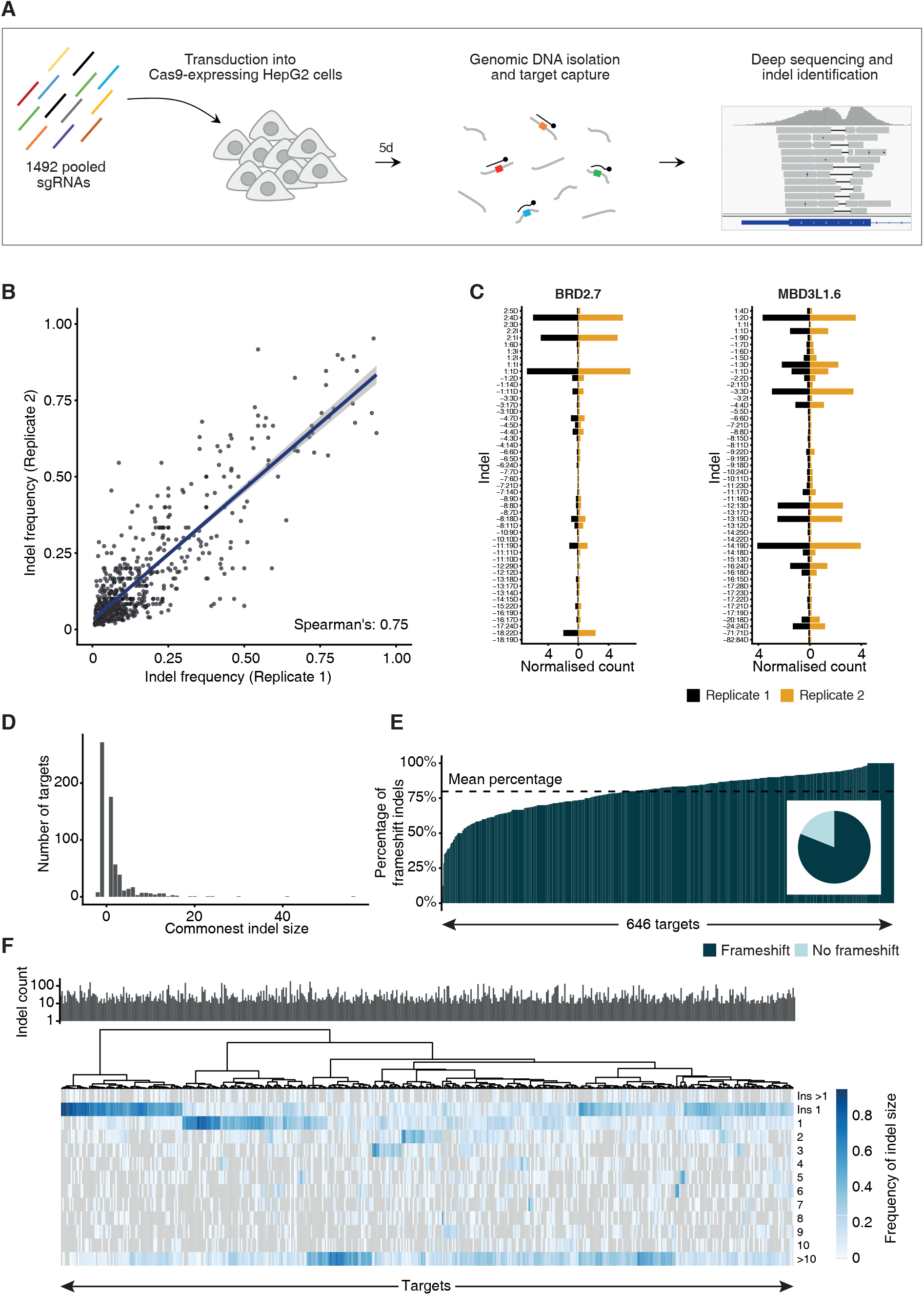
General specificity and reproducibility of CRISPR-mediated indel profiles at individual sites. **A**. An overview of the experimental setup. 1492 pooled sgRNAs were transduced into Cas9 expressing HepG2 cells. Targeted capture of DNA around cleavage sites was followed by deep sequencing and indel identification. **B**. CRISPR/Cas9 cleavage results in highly reproducible indels at over 1000 target sites. The frequency at which each detected indel occurs at each target site is compared across two biological replicates. **C**. Indel profiles are highly reproducible at individual target sites. Validation with deeper sequencing shows the indel profiles for two biological replicates at the BRD2.7 and MBD3L1.6 target sites. The indel nomenclature is [start coordinate relative to cleavage site]:[size][insertion or deletion]. Counts are normalised to the total library size for each experiment. **D**. Indels are commonly mononucleotide insertions or deletions. The histogram shows the size of the commonest indel for each target. **E**. Indels usually result in frameshift mutations. For each target, the percentage of indels resulting in a frameshift mutation is shown. The inset pie chart shows the proportion of targets for which the commonest observed indel is a frameshift mutation. **F**. CRISPR-mediated editing results in diverse patterns of indels. The heatmap shows the frequency at which indels of a given size occur at each target site. The target sites are clustered using Ward D2 hierarchical clustering. The bar plot above indicates the number of indels observed at each target site below. Only data from targets from the 450 pools (525 targets) are used to enable fair comparisons.

In agreement with previous studies that examined a limited number of sites (Brinkman et al., 2014; van Overbeek et al., 2016), we observed that RGN-induced editing is highly reproducible across biological replicates (Spearman’s coefficient 0.75, p < 2.2 × 10^−16^) indicating that repair outcome is non-random (Fig. 1B). High-coverage analysis of cells transduced with individual sgRNAs confirmed these results showing almost identical indel patterns in two independent experiments (Fig. 1C). Furthermore, our ability to probe a large number of sites simultaneously allowed us to reveal general patterns of CRISPR-mediated DNA editing and make a number of observations. First, single-nucleotide indels were the most frequent type of indel for the majority of targets, with 42% and 27% of targets showing 1 nt insertions or deletions respectively as their commonest indel (Fig. 1D). Nevertheless, sites showing a preference for longer deletions, of up to 56 nt, were also observed (Fig. 1D). Second, in line with the observed bias for single-nucleotide alterations, CRISPR-induced indels often resulted in frameshift alterations (Fig. 1E). On average, 79.8% of indels induced at a given site disrupted the gene coding frame, a percentage significantly higher than the theoretical 66% assuming a random outcome (p < 2.2 × 10^−16^, χ^2^ test) (Fig. 1E). Moreover, 89% of all detected indels resulted in a frameshift (Fig. 1E). Thus, the probability of achieving protein loss-of-function through CRISPR-induced indels is typically relatively high. However, 3 sites showed strong preference for in-frame indels (in-frame indels ≥ 70%), suggesting that in certain cases it may be difficult to successfully induce gene KO. Third, unsupervised hierarchical clustering identified four groups of targets showing similar indel patterns (Fig. 1F). Based on the relative frequency of the observed indels, targets could be broadly divided into sites that preferentially show small insertions, small deletions, long deletions or have no clear preference (Fig. 1F). Fourth, sgRNA activity, as measured by quantifying indel counts at each site, was highly variable, ranging from 0 to 188 (Fig. S2C and Fig. 1F). Indel count did not correlate with abundance of sgRNAs in the pools, suggesting that sgRNA activity is intrinsically variable (Fig. S2D) (R^2^ = 0.02). This observation is in agreement with previous findings obtained by inferring sgRNA activity from their ability to induce an expected phenotype (Doench et al., 2016; Wang et al., 2014). Of note, several inactive sgRNAs had high predicted activity scores, indicating that predicting algorithms can be further improved, and that in addition to DNA sequence, other factors may affect sgRNA activity at a given site (Fig. S1B). Activity did not correlate with preference for a certain type of indel pattern (Fig. 1F).

### Precision of CRISPR-induced DNA editing varies considerably across sites

The observation that different targets display distinct preferences for certain indel types prompted us to examine the degree of precision of DNA editing, i.e. the preference for a specific indel, across sites. To do so, we first calculated the relative frequency of each distinct indel, defined by its coordinates and base composition, at each site and then ranked all sites based on the frequency of the commonest indel. This analysis revealed a large range of precision, with some targets displaying up to 79 distinct, infrequent indels (frequency < 5%) and others showing one dominant indel (up to 94% frequency) and only a few additional ones (Fig. 2A – D). Overall, we found that for approximately one fifth of the targets there is at least a 50% chance of inducing a specific indel, but the majority of sites are more unpredictable. On average, the commonest indel frequency for a given site was 33.9% and the median number of observed distinct indels was 12.

**Figure 2.**
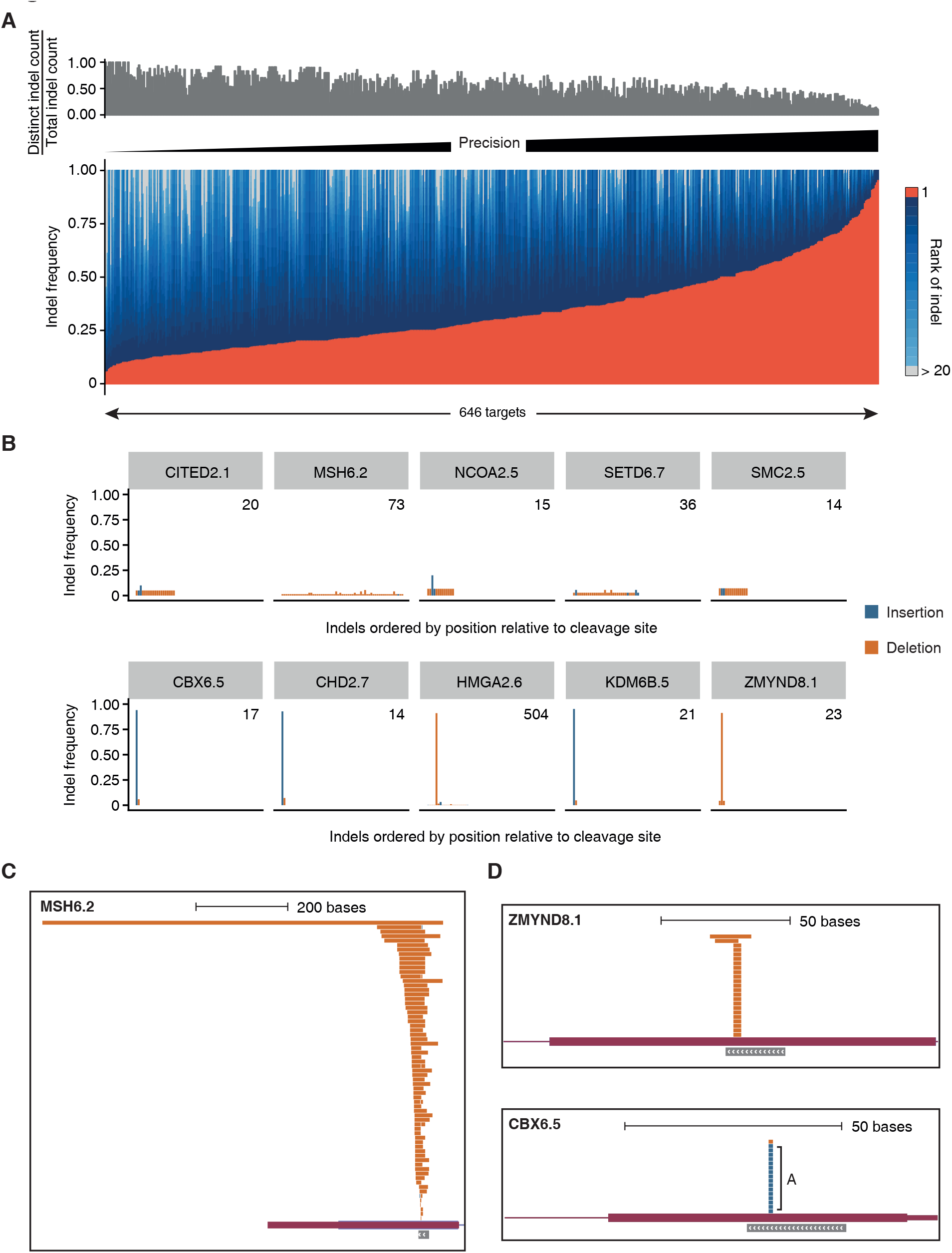
Site-specific editing precision. **A**. The commonest indel detected at a site has a wide range of frequencies. For each of the 646 target sites the frequency of each indel is shown. The most common is coloured red, while the subsequent ranks of indel are progressively lighter shades of blue, with those ranked higher than 20 in grey. The bar plot above shows the number of distinct indels relative to the total number of detected indels at each site. **B**. Indel profiles of the 5 most imprecise (above) and the 5 most precise (bottom) targets. Along the x-axis, the indels are ordered by start coordinate relative to the cleavage site. The inset number indicates the total number of indels detected at that site. **C**. Example of a site showing a wide range of distinct indels. The indel profile for MSH6.2 is shown in orange, with the gene body coloured in plum and the sgRNA binding position in grey, with chevrons indicating strand. **D**. Examples of sites showing strong preference for a specific deletion (ZMYND8.1) or insertion (CBX6.5). Note that all insertions at CBX6.5 show introduction of the same nucleotide, A.

### Editing precision correlates with editing efficiency, indel type and indel size

To examine the relationship between editing precision and indel features, we categorized target sites into three groups: imprecise (0 < commonest indel frequency ≤ 0.25), medium (0.25 < commonest indel’s frequency ≤ 0.5) and precise sites (0.5 < commonest indel frequency ≤ 1), with each group containing comparable numbers of sites (Fig. 3A). Notably, editing precision correlated with efficiency of indel formation (p < 2.2 × 10^−16^, Kruskal-Wallis test) (Fig. 3B). Precise targets showed on average twice as many indels as imprecise targets, and the most active sites showed a strong preference for specific indels (commonest indel’s frequency > 0.57) (Table S2). We then asked whether editing precision correlated with preference for insertions or deletions. Imprecise targets showed a high proportion of deletions, with insertions being on average only 20% of the total indels, whereas insertions were more frequent in the middle group of targets (Fig. 3C). Precise targets segregated into two distinct subsets: 67.6% showed a strong preference for insertions whereas the rest mainly repaired RGN-induced breaks by inducing deletions (Fig. 3C). The two subsets were clearly separated, likely reflecting their tendency to induce mainly one dominant indel. Editing precision also correlated with absolute indel size (Fig. 3D). While imprecise and middle targets showed a range of indel sizes, with deletions as long as 2315 bp, precise targets displayed a strong bias towards single-nucleotide indels (Fig. 2B, 3D, E). Combining insertion and deletions, 71% of edited sequences in the precise group had a single-nucleotide alteration. We conclude that RGN-based editing precision varies considerably across sites and correlates with editing efficiency and with the type of resulting indels.

**Figure 3.**
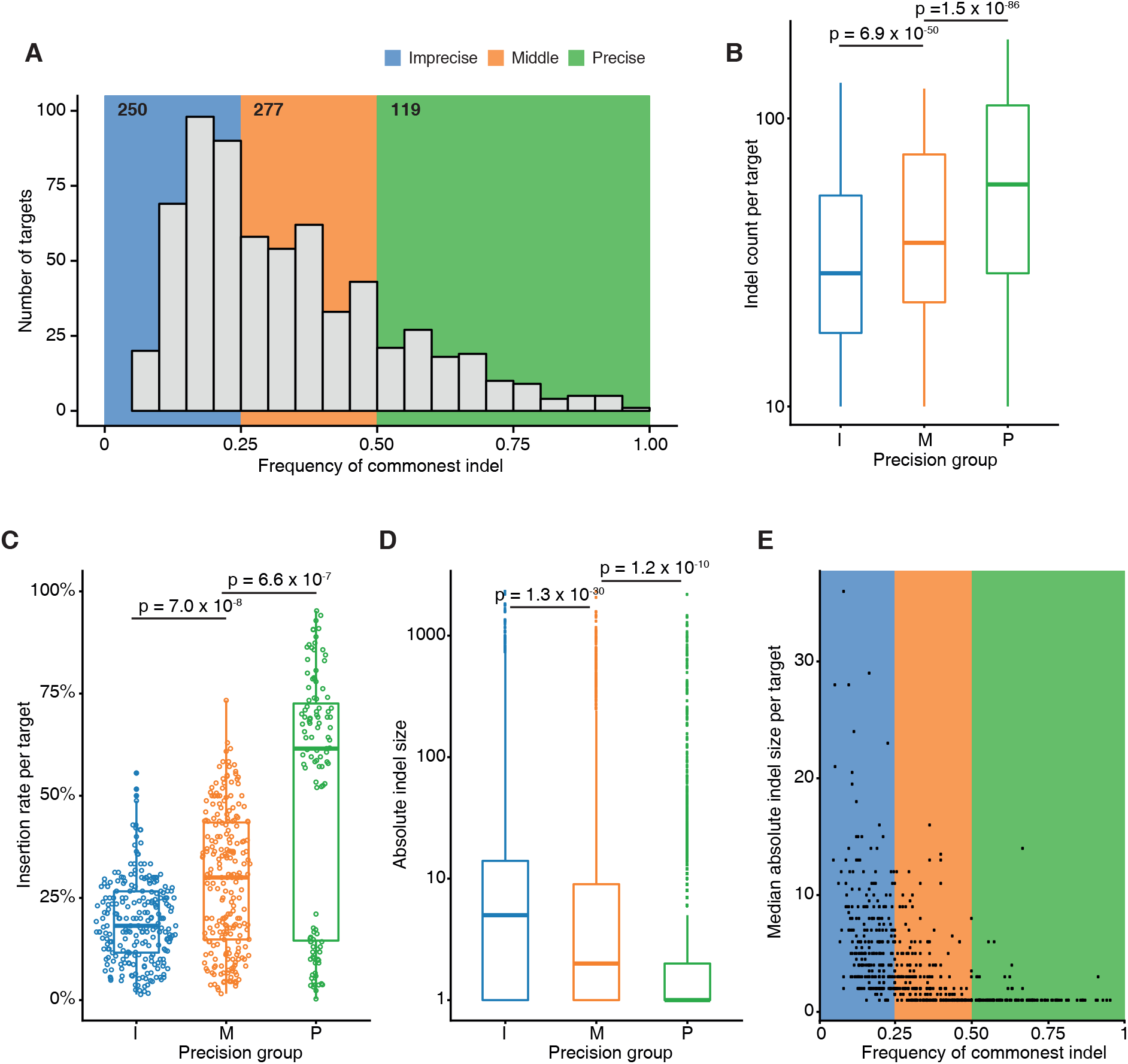
Relationship between editing precision and indel features. **A**. The distribution of commonest indel frequencies at target sites is shown. The background indicates three groups of sites as defined based on their editing precision. Inset numbers indicate the number of target sites in that group. **B**. Efficiency of indel formation increases with precision of the target site. The indel count for each target site in a group is shown, with only data from the 450 pools used to enable fair comparisons. I: imprecise, M: middle, P: precise. Statistical testing was performed using the Kruskal-Wallis test followed by Dunn’s test for multiple comparisons with Benjamini-Hochberg correction for multiple testing. **C**. Precise targets can be further categorised into insertion-preferred and deletion-preferred. The percentage of indels that are insertions at each target are shown, grouped by precision. Statistical testing was performed using the Kruskal-Wallis test followed by Dunn’s test for multiple comparisons with Benjamini-Hochberg correction for multiple testing. **D**-**E**. Precise targets have shorter indels. The mean absolute indel size for each of the three groups is shown in D. The median absolute indel size is compared with the frequency of the commonest indel (i.e. the measure of precision) at each target site in E. The background is coloured as in A. Statistical testing was performed using the Kruskal-Wallis test followed by Dunn’s test for multiple comparisons with Benjamini-Hochberg correction for multiple testing.

### Precise targets exhibit primarily homology-associated insertions and deletions

Although indel profiles have been shown to be dependent on both MH-dependent and MH-independent mechanisms (Bae et al., 2014; Brinkman et al., 2018; van Overbeek et al., 2016), a quantitative assessment of their relative contribution across many target sites is lacking. In the absence of genetic or pharmacological interference with specific repair pathways (e.g. NHEJ, HDR or MMEJ), characterisation of indel profiles is insufficient to determine which specific mechanism led to an observed outcome. We therefore performed a pathway-agnostic analysis of indels that searched for any homology at the indel boundaries. This analysis revealed that MH of variable size, ranging from 1 to 18 nt, characterised the majority of deletions (Fig. 4 A-C, Table S3). 73.1% of all deletions showed evidence of MH-mediated repair (MH-deletions) and on average, 75% of deletions at a given site were characterized by MH (Fig. 4A). Deletions associated with shorter MHs (1-4nt) were also enriched above the expected frequency, indicating that the effect of sequence homology on repair outcome is not limited to longer MH stretches (5-25nt) used by the MHEJ pathway (Fig. 4B). MH-deletions were enriched in the groups of precise and middle targets (p = 3.2 × 10^−6^, Kruskal-Wallis test) (Fig. 4D). Furthermore, regardless of editing precision, 80% of targets had a MH-deletion as their commonest one.

**Figure 4.**
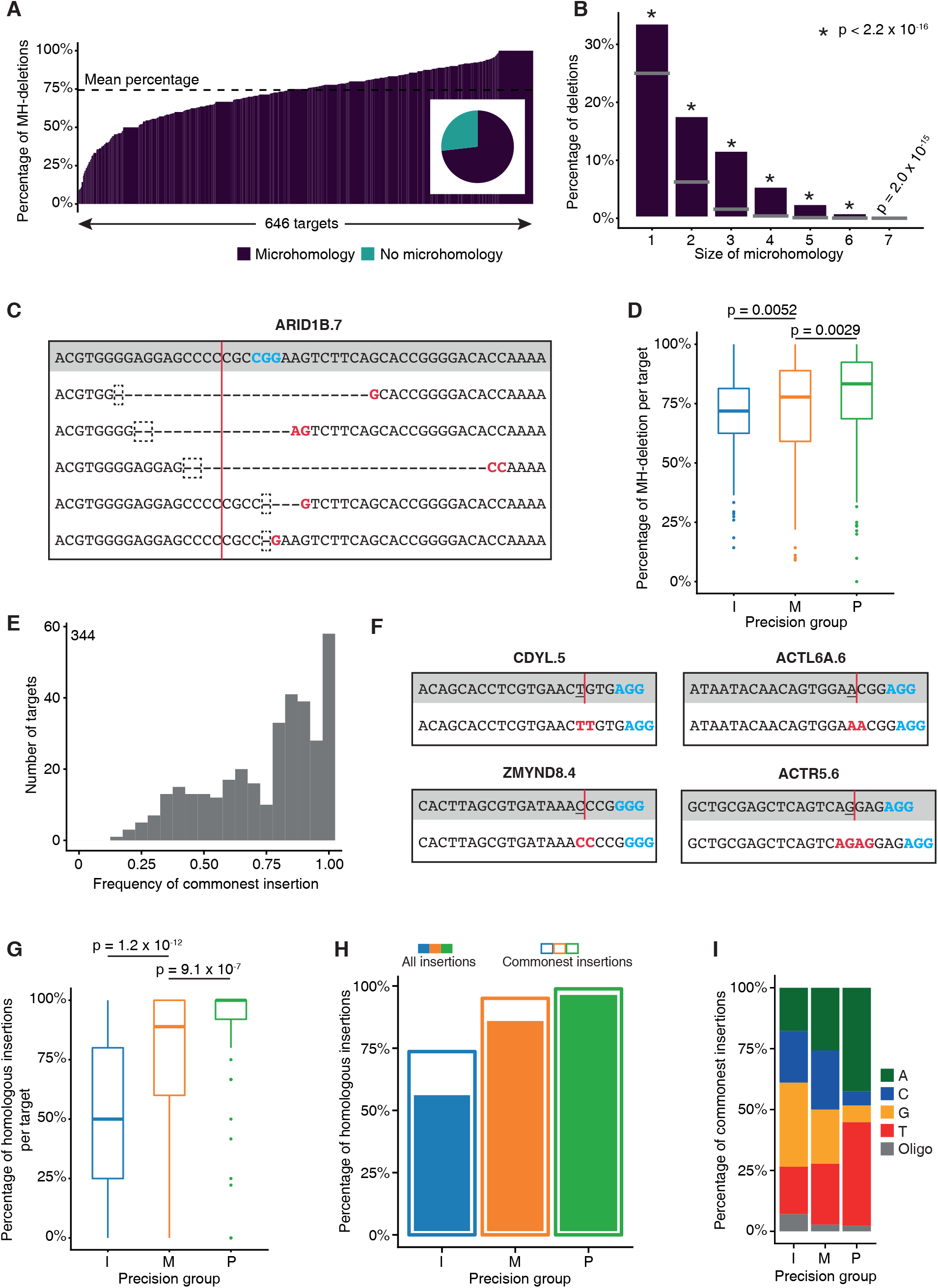
Precise targets are enriched for homology-associated indels. **A**. The majority of deletions show evidence of microhomology-guided end-joining. The percentage of microhomology-associated deletions (MH-deletions) at each target site is shown. In the inset pie-chart, the proportion of all detected MH-deletions is shown. **B**. A range of microhomology sizes is observed. The percentage of deletions that have microhomology of a given size is shown. The grey bar indicates the expected percentage for each *k*-mer size. Statistical testing was performed using the chi-squared test. **C**. An example of microhomology-associated repair. All the deletions at the ARID1D.7 target site are shown. Above in the grey panel is the reference sequence, with the protospacer adjacent motif emboldened in blue and the expected cleavage site indicated with a red line. Below, each line represents each distinct deletion detected. In the dashed box is the microhomology in the deletion and emboldened in red is the corresponding microhomology in the unedited part of the sequence. **D**. Precise targets are enriched for microhomology-associated deletions. The percentage of MH-deletions at individual target sites grouped by precision are shown. I: imprecise, M: middle, P: precise. Statistical testing was performed using the Kruskal-Wallis test followed by Dunn’s test for multiple comparisons with Benjamini-Hochberg correction for multiple testing. **E**. Recurrent insertions upon CRISPR-mediated editing. The frequency of the commonest insertion at a target site is shown. Only targets with 5 or more insertions are considered to obviate a low-count bias. The inset count is the number of target sites included. **F**. Examples of insertion homology. Insertions are shown for the CDYL.5, ACTL6A.6, ZMYND8.4 and ACTR5.6 target sites. Above in the grey panel is the reference sequence, with the protospacer adjacent motif emboldened in blue and the expected cleavage site indicated with a red line. The −4 position is underlined. Below, the edited sequence is shown with the insertion homology emboldened in red. For the first three this is a mononucleotide; for ACTR5.6, a dinucleotide. **G**. Precise targets are enriched for homologous insertions. The percentage of homologous insertions at individual target sites grouped by precision are shown. Statistical testing was performed using the Kruskal-Wallis test followed by Dunn’s test for multiple comparisons with Benjamini-Hochberg correction for multiple testing. **H**. Commonest insertions often exhibit insertion homology. The percentage of all homologous insertions in a group is shown in filled bars. In the outlined bars is the corresponding percentage of commonest insertions. **I**. Precise targets often insert adenosine and thymine nucleotides. The nucleotide inserted as the commonest insertion for each group is shown.

Although sequence homology has not been implicated in the formation of insertions, surprisingly we found that many target sites showed recurrent insertions containing a common inserted base, suggesting that the choice of inserted nucleotide is non-random (Fig. 2D, Fig. 4E and Table S4). Moreover, the recurrently inserted base was often homologous to nucleotide −4 from the PAM sequence, which is typically the nucleotide upstream of the cleavage site (Jinek et al., 2012) (89% of the commonest insertions at each target) (Fig. 4F); we termed this feature “insertion homology”. As observed for deletions, the prevalence of insertion homology correlated with editing precision (p < 2.6 × 10^−16^, Kruskal-Wallis test) (Fig. 4G,H). Precise targets displayed 96% of homologous insertions, whereas this percentage was only 56% in the imprecise group (p < 2.6 × 10^−16^, χ^2^ test) (Fig. 4H), suggesting that template-mediated insertions are a strong determinant of the observed site-specific indel profiles. Even at imprecise targets, homologous insertions were often the commonest ones (Fig. 4H). Notably, precise targets showed a strong bias for inserted “A”s and “T”s, suggesting that sequence features underlie the correlation between editing precision and homologous insertions (Fig. 4I). Altogether, these observations suggest that homology-mediated end-joining strongly influences DNA repair outcome, both for insertions and deletions, and correlates with site-specific precision of CRISPR-mediated editing.

### DNA sequence determines editing precision in a predictable manner

To examine whether editing precision depends on the base composition of target sites, we employed a machine learning approach to identify features that distinguish precise and imprecise target sites. We trained neural networks that predict precision, i.e. commonest indel frequency. We bootstrapped with replacement 10 times using 80% of the targets selected randomly to train the networks, with the remaining 20% kept unseen for testing each time. We found a significant correlation between the predicted and observed precisions for the 129 test target sites (correlation coefficient R^2^ = 0.40-0.57, p-value =2.31×10^−12^-2.45×10^−6^, Wald tests) (Fig. 5A, B). Although the predictive power of the model was only moderate, it allowed us to identify important positions in the protospacer. If certain positions have a significant influence on precision prediction, randomising those nucleotides should dramatically reduce the correlation with observed precision. To investigate this, we performed a permutation “nucleotide” importance analysis, systematically randomising each position in test sequences and examining the resulting effect on their predicted commonest indel frequency. This analysis revealed that the nucleotide at position −4 from the PAM sequence had the strongest influence on precision as a single nucleotide, reducing the predictive power by 30% when randomised (average R^2^ = 0.36) (Fig. 5C). Nucleotide positions −2, −3, and −5 also showed an effect, although weaker, reducing R^2^ by 12%, 5% and 14% respectively. Simultaneous randomisation of all 4 of these nucleotides reduced R^2^ by over 80% and abolished the predictive significance of the trained model (average R2 = 0.10 and average p = 0.356, Wald tests), indicating that these positions within the protospacer, especially the one upstream of the cleavage site, are critical for defining editing precision at a target site (Fig. 5D). We refer to these combined nucleotides as the “precision core” of a target site.

**Figure 5.**
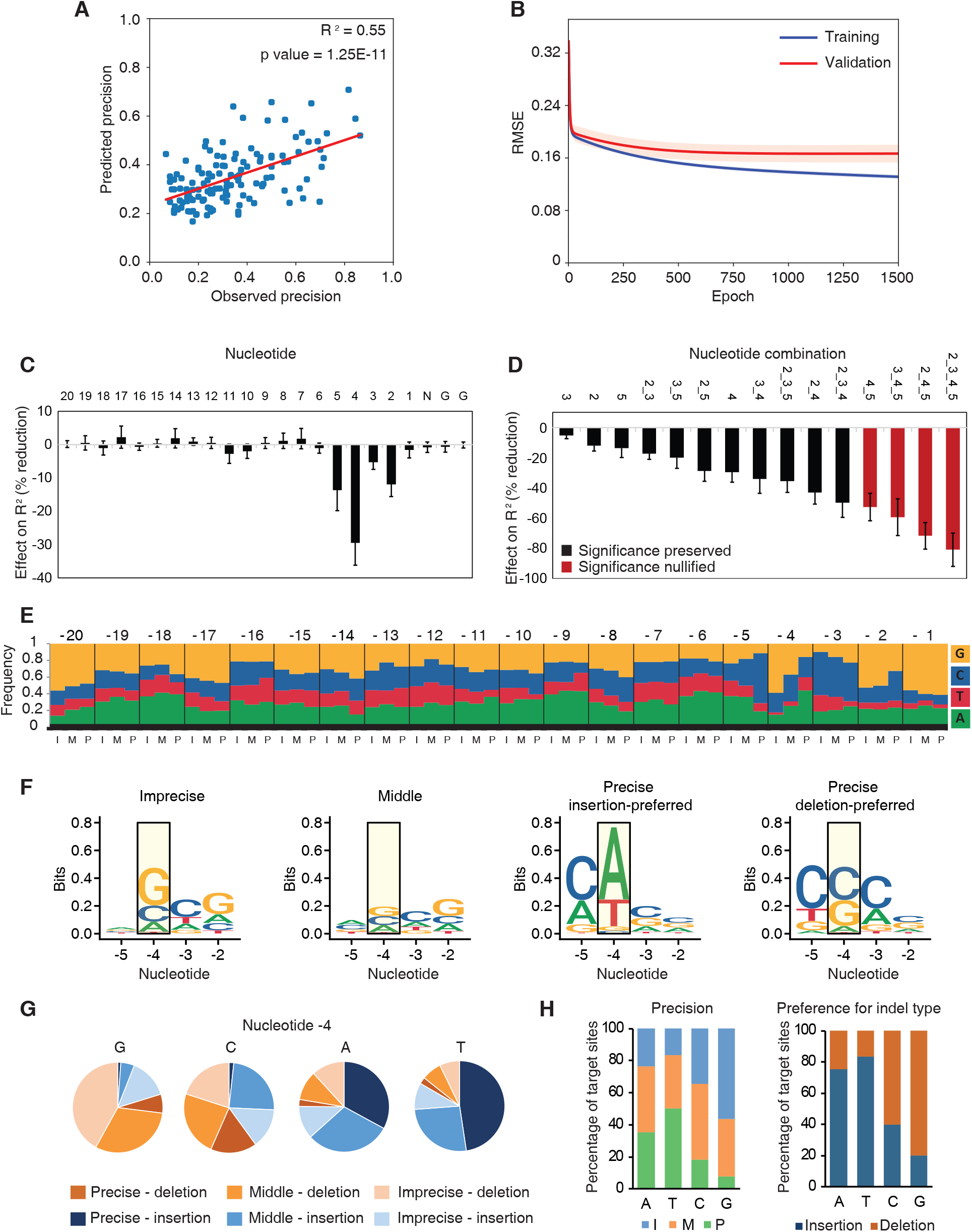
A neural network identifies protospacer nucleotide positions that are important for determining precision. **A**. Correlation between the observed precision at a given target site and that predicted by the neural network. One of ten trained neural network models is shown (see Methods). The coefficient of determination is shown, with statistical significance testing using Wald χ^2^ test. **B**. Training the artificial neural network using random subsampling resulted in consistent, statistically significant, predictive capability. Error margins (orange shade) are standard deviation from the mean RMSE (root-mean squared error of predictions vs observed values) obtained from training the neural network with 10 random sgRNA subsamples. Epoch indicates the number of iterations the network was presented with the training dataset. **C**-**D**. Contribution of the indicated protospacer nucleotides (C) or combination of nucleotides (D) to editing precision. The effect of nucleotide randomization is shown as reduction of the model’s predictive power (R^2^). Values are mean and standard deviation from 10 different trained neural networks. Bars in red indicate randomized positions that increased the average p value of Wald tests across models to nonsignificant levels (p >0.05). **E**. The frequencies of each nucleotide in the protospacer sequence for each precision group is shown. I: imprecise, M: middle, P: precise. **F**. Sequence logos for the combined 4 key positions of the protospacer for the different precision groups are shown. Precise targets are split based on their preference (commonest indel) for insertions or deletions. The most important −4 nucleotide position is highlighted in a yellow box. **G**. The proportion of target site groups that have the indicated nucleotide at the −4 position is shown. Sites are grouped based on their precision and their preference (commonest indel) for insertions or deletions. **H**. The percentage of target site groups that have the indicated nucleotide in the −4 position is shown. Sites are grouped based on their precision (left) or their preference (commonest indel) for insertions or deletions (right).

Examination of base preference throughout the protospacer sequence revealed significant differences at many positions when comparing targets of different precision groups (Fig. 5E). In agreement, different groups displayed distinct sequence motifs at the precision core, with the middle group showing the least defined motif, as expected (Fig. 5F). Notably, precise targets showed distinct base preference depending on whether their commonest indel was an insertion or a deletion (Fig. 5F). As expected, nucleotide −4 showed the biggest differences, followed by nucleotide −5, which was frequently a “C” specifically in precise targets (Fig. 5F). We then examined to what extent nucleotide −4 on its own could predict editing outcome. Different bases at position −4 showed distinct association with indel types (insertions vs deletions) and precision groups (Fig. 5G). The vast majority of target sites that contained an “A” or a “T” upstream of the cleavage site repaired RGN-induced DSBs via insertions (75% and 83% of targets, respectively) (Fig. 5H, Fig. S3). These were mostly precise or middle targets (median commonest indel frequency: 0.41 and 0.48 for targets with “A” and “T”, respectively) (Fig. 5H, Fig. S3). When taking into account positions −5 and −4 together, the correlation with precision further increased (median commonest indel frequency: 0.53 and 0.65 for targets with “CA” and “AT”, respectively) (Fig. 5F and Fig. S3). In contrast, 80% of targets containing a “G” at position −4 showed deletions and were mostly imprecise targets (median commonest indel frequency: 0.22) (Fig. 5H, Fig. S3). Moreover, 76% of targets containing “CC” at positions −5 and −4 induced relatively precise deletions (median commonest indel frequency: 0.40) (Fig. 5F and Fig. S3). Given the large number of sites examined, the observed percentages assume a predictive value with respect to the editing outcome that may occur at similar protospacers. We conclude that CRISPR-induced editing precision at a given site can be predicted by examining the base composition of the precision core, and that position −4 is sufficient to predict with a high degree of confidence whether a site will repair by introducing insertions or deletions.

### Chromatin states affect RGN activity

Our findings, in agreement with previous small-scale studies (Brinkman et al., 2014; van Overbeek et al., 2016), suggest that DNA sequence features strongly affect RGN-induced indel profiles in a site-specific manner, influencing editing precision and efficiency. However, even within precision groups, the number of induced indels and their patterns varied across sites (Fig. 3B). Furthermore, the predictive power of the neural network model, based solely on the protospacer sequence, was moderate, suggesting that other factors may influence indel profiles. We therefore examined whether chromatin structure may contribute to the observed editing outcome. To do so, we selected 6 target sites characterized by variable editing precision and efficiency of indel formation (Fig. 6A), and individually transduced the corresponding sgRNAs in Cas9-expressing cells in the presence of chromatin-modulating compounds. We used the HDAC inhibitor Trichostatin A (TSA) to induce global histone hyperacetylation, and the EZH2 inhibitor GSK126 (EZH2i) to reduce the levels of H3K27me3 (Fig. S4A). Both treatments significantly affected the efficiency of indel formation inducing dose-response changes (p < 0.001, paired Wilcoxon test) but while TSA increased the number of detected indels, reaching almost a 2-fold increase for the ACTL6A.5 site, EZH2i had the opposite effect and inhibited indel formation (Fig. 6B). The effect was highly reproducible across biological replicates (Fig. S4B,C Table S5). The effect of TSA varied depending on the target and inversely correlated with the endogenous levels of histone modifications (Fig. 6B,C). Sites characterized by low levels of acetylated H3 showed a greater response to the treatment compared with those that already had high levels of the endogenous marks (MSH6.2 and SMARCD2.1), suggesting a direct effect of chromatin modulation on indel formation (Fig. 6B,C). Analysis of individual indels indicated that the effect of TSA and EZH2i was not restricted to a few indels and that both insertions and deletions were affected (Fig. 6D, Fig. S5A, Table S5). We conclude that the chromatin state of target sites affects the activity of RGNs, and that induction of histone acetylation enhances DNA editing efficiency.

**Figure 6.**
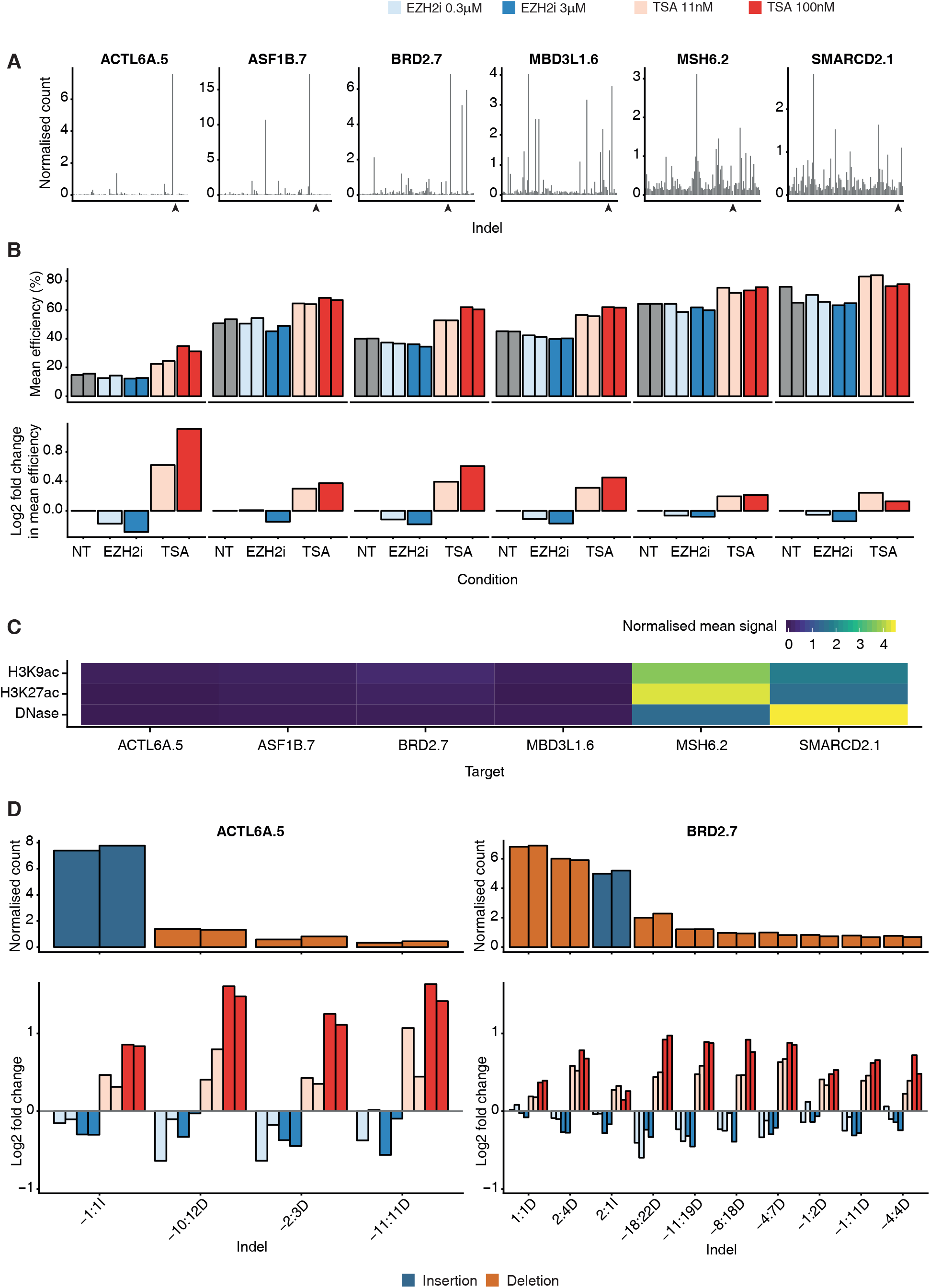
Chromatin modulation affects RGN activity. **A**. The indel profile at each of the selected target sites in untreated cells. Along the x-axis, the indels are ordered by start coordinate relative to the cleavage site (indicated with an arrowhead), with the count of each distinct indel on the y-axis, normalised by the effective library size at that site. The mean across both replicates is shown. **B**. Mutation efficiency is altered upon chromatin modulation in a dose-responsive manner. Above is shown the mutation efficiency for each of the treatments for both replicates separately. Below is shown the log_2_ fold-change in mean efficiency relatively to untreated cells (NT) for each corresponding target site and condition above. **C**. Mean ChIP-seq signal for H3K9ac and H3K27ac and DNase-seq signal in untreated HepG2 cells. Signal in a 500-nucleotide window centred on the cleavage site at each target site is shown as a heatmap. **D**. Chromatin modulation affects both insertions and deletions ACTL6A.5 and BRD2.7 are shown as examples (see also Fig. S5). Above is shown the count of each indel, normalised by the effective library size at each site for each replicate. Only indels with a normalised count of at least 1 in any condition are included. The indel nomenclature is [start coordinate relative to cleavage site]:[size][insertion or deletion].

### Chromatin states influence indel profiles

Although changes in editing efficiency by TSA or EZH2i were observed for most indels at each site, some indels were preferentially affected (Fig. 6D). Furthermore, shorter and longer indels appeared differentially affected by treatment (Fig. S5B). These observations suggest that chromatin modulation may alter indel profiles. We therefore examined the relative changes in the abundance of individual indels, focusing on the effect of TSA, which induced greater and more consistent changes in indel formation (Fig. 6B,D). Across all sites, we observed dose-dependent changes in the relative frequency of indels, with some being favoured at the expense of others (Fig. 7A,B, Fig. S6). Although the overall indel patterns were maintained, confirming robustness of the editing profiles, the most frequent indels showed reproducible and dose-dependent changes (Fig. 7A-C). At some sites (MBD3L1.6, MSH6.2 and SMARCD2.1) the preference for their commonest indel was enhanced, while at others (ACTL6A.5, ASF1B.7 and BRD2.7) it was decreased (Fig. 7 A-C). Notably, at the BRD2 site, the identity of the commonest indel changed upon TSA treatment, suggesting that changes in chromatin structure can alter indel profiles (Fig. 7 A-C). Altogether, these results show that chromatin structure contributes to the establishment of site-specific indel profiles. While DNA sequence appears to be the major determinant of CRISPR-mediated editing outcome, the chromatin state of a given site modulates the relative abundance of individual indels and contributes to defining the site’s indel profile. The combination of genetic and epigenetic factors makes site-specific patterns highly robust and reproducible.

**Figure 7.**
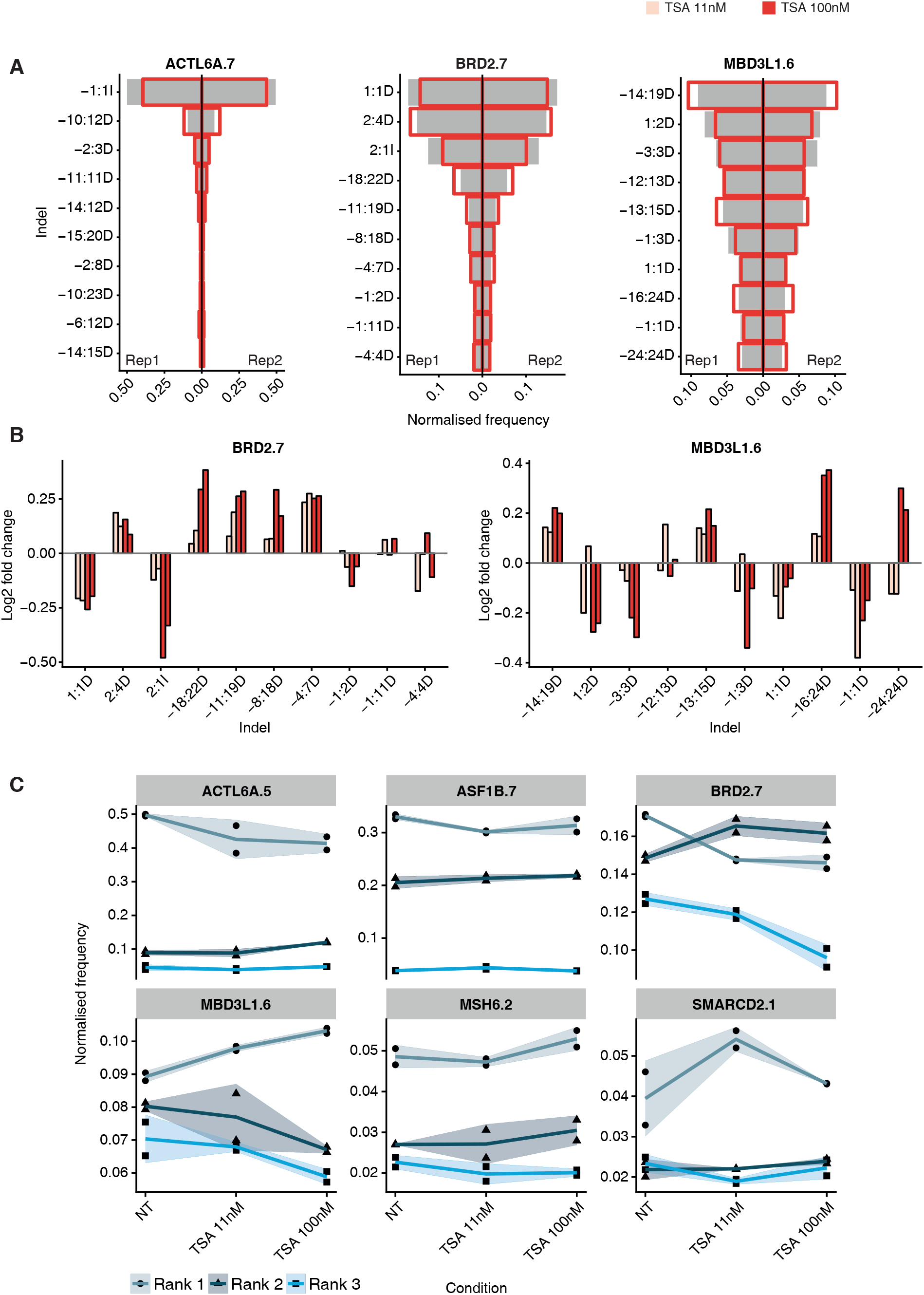
Chromatin modulation affects indel profiles. **A**. Chromatin modulation has a differential effect on distinct indels. ACTL6A.7, BRD2.7 and MBD3L1.6 are shown as examples (see also Fig. S6). The count for each indel, normalised by the total number of indels detected at that target site in that condition is shown for each replicate. The frequency of the indicated indels in the untreated condition (grey bars) and in the TSA 100nM condition (red outline) is shown. The indel nomenclature is [start coordinate relative to cleavage site]:[size][insertion or deletion]. The 10 commonest indels for each site are shown. **B**. The log_2_ fold change in the frequency of the indicated indels is shown for BRD2.7 and MBD3L1.6 (see also Fig. S6). The 10 commonest indels across both replicates are shown. **C**. The change in frequency is shown for the 3 commonest indels (ranks 1, 2, 3) for all 6 validated target sites. The line indicates the mean of both replicates, the shaded area is mean ± 1 standard deviation. NT indicates untreated cells.

## DISCUSSION

### Predicting editing precision

Although the bacterial CRISPR system has been widely adopted as the preferred genome engineering tool, our ability to predict editing accuracy, efficacy and outcome at specific sites is still limited. A major obstacle in defining precise genome editing rules is our incomplete understanding of how RGNs interact with eukaryotic cellular components – complex genomes containing repetitive sequences, the packaging of DNA into chromatin and the presence of various cellular pathways that recognize and repair RGN-induced DSBs. Various studies have provided insights into some of these interactions (Brinkman et al., 2018; Isaac et al., 2016; Jensen et al., 2017; Kosicki et al., 2018; Lemos et al., 2018; van Overbeek et al., 2016). However, due to the limited number of characterized target sites, discerning whether the observed patterns are general or site-specific features is not straightforward. Through systematic analysis of indel formation at over 1000 different sites in the human genome, this study reveals general trends of CRISPR editing and provides simple rules to predict how a given target may respond to RGN-induced DSBs.

Extending the observation that indel profiles are non-random (van Overbeek et al., 2016), we find that precision of DNA editing varies considerably among sites, with some targets showing one highly-preferred sequence alteration and others displaying a wide range of infrequent, yet reproducible, indels. We show that editing precision is an intrinsic feature of the target site and depends on four nucleotides located around the cleavage site within the protospacer, with the most influential position being the −4 nucleotide from the PAM sequence. Strikingly, the mere presence of a “T” at position −4 gives a site a 50% probability of repairing in a precise manner. Our finding that editing precision is site-specific and can be predicted has important implications. For practical reasons, knowing what editing outcome is likely to occur at a given site has obvious advantages and maximizes the chance of having a desired sequence alteration, both for clinical and research applications. Although pharmacological modulation of repair pathways has been shown to modulate indel profiles, the induced changes are subtle, and for many applications use of inhibitors may not be suitable (van Overbeek et al., 2016; Shou et al., 2018). Targeting a likely precise site would be a more effective way of steering CRISPR-mediated editing towards a desired outcome. Moreover, given the extreme reproducibility of indel patterns, prediction of a precise target combined with experimental validation in model systems could considerably increase safety of clinical editing applications. This is particularly relevant in light of recent studies reporting the occurrence of large on-target deletions that may have pathological consequences (Kosicki et al., 2018).

### Relationship between editing precision and indel type

Our findings also reveal a strong correlation between editing precision and preference for repairing RGN-induced DSBs via insertions. We show that targets with “A”s or “T”s at nucleotide −4 mainly show insertions, with the commonest insertion being highly recurrent and representing on average approximately half of the indels detected at a given site (Fig. 5H, Fig. S3 and Table S4). As we write, Taheri-Ghahfarokhi et al. have reported similar findings upon analysis of 15 target sites (Taheri-Ghahfarokhi et al., 2018). The identity of the recurrent insertions can also be predicted, as the inserted nucleotide is nearly always homologous to the −4 nucleotide (Fig. 4G-I and Table S4). Such predictions could, for instance, allow efficient induction of a stop codon (TAA) when an in-frame TA dinucleotide is present at positions −5 and −4 of the targeted region. In contrast, targets with “G”s at nucleotide −4 are the most imprecise and mainly repair inducing a variety of unpredictable deletions (Fig. 5H and Fig. S3). Thus, choosing target sites with “A”s or “T”s at nucleotide −4 is an effective way to induce predictable insertions at a regions of interest.

### Critical role of nucleotide −4 in defining site-specific indel profiles

The key role of nucleotide −4 in influencing editing precision and preference for indel type is particularly interesting in light of recent findings that revealed flexible scissile profiles by Cas9 and generation of 5’ overhangs upstream of the canonical cleavage site, due to asymmetric cleavage of the two DNA strands (Shou et al., 2018). Notably, 5’ overhangs are mostly observed at position −4 on the non-complementary strand. These findings, together with our results, explain the prevalence of single-nucleotide insertions homologous to the −4 nucleotide, as the overhanging nucleotide can be used as a template before ends are re-joined. Thus, paradoxically, imprecision of Cas9 cleavage is the likely cause of precision in the insertion outcome. Similarly, the high frequency of single-nucleotide deletions is likely related to the asymmetric cleavage of DNA by Cas9.

Envisioning how the base composition of position −4 may influence editing precision is not straightforward. One possibility is that the nature of the 5’ overhanging nucleotide may recruit distinct proteins involved in DNA repair. Alternatively, it may affect Cas9 binding to the broken ends and this may, in turn, affect the repair outcome. The other nucleotides in the precision core may act similarly. Structural analysis of RGNs with distinct −4 nucleotides may help shed light on this issue.

Our observation that the vast majority of detected insertions show homology, combined with the finding that NHEJ-mediated repair of CRISPR-induced DSBs is mostly error-free (Geisinger et al., 2016), and that deletions generated by sgRNA pairs can be repaired with a high level of precision (Shou et al., 2018), suggest a model whereby flexible cleavage by Cas9 influences DNA repair fidelity: when blunt ends are generated at nucleotide −3, cells repair DSBs in an error-free manner reconstituting the original sequence, whereas indels occur mainly when asymmetric cleavage generates overhanging ends. This model may also reconcile apparently conflicting results about the fidelity of NHEJ in CRISPR-independent and CRISPR-dependent contexts (Brinkman et al., 2018; Dudley et al., 2005; Geisinger et al., 2016; van Heemst et al., 2004; Shou et al., 2018). Interestingly, both outcomes are useful for genome editing, as blunt ends allow precise genomic deletions and insertions of exogenous sequences, while overhanging ends enable induction of indels resulting in gene KO.

### Influence of chromatin environment on site-specific editing outcome

Although DNA sequence is a major determinant of site-specific indel profiles, we show that packaging of DNA into chromatin affects editing efficiency and the relative frequency of indels at a given locus. We find that histone hyperacetylation and reduction of the heterochromatin-associated mark H3K27Me3 induces opposite changes in editing efficiency, enhancing and inhibiting indel formation, respectively. Although the effect of TSA was observed at all tested sites, the effect was particularly pronounced at sites with low endogenous levels of histone acetylation, suggesting that transient TSA treatment may be a strategy to enhance editing efficiency at sites located in repressive chromatin environments. Our results are in agreement with the observed correlations between sgRNA activity and open chromatin at the genome-wide levels and evidence from *in vitro* studies indicating that nucleosome positioning impairs binding of Cas9 to DNA and inhibits its activity (Fig. 6C) (Horlbeck et al., 2016; Uusi-Mäkelä et al., 2018). In addition to interfering with Cas9 binding to a target site, chromatin may also affect its cleavage profile, favouring either blunt ends that can be precisely repaired or overhanging ends that promote the formation of indels. We also show that modulation of chromatin differentially affects individual indels at a target site, and can change the identity of the commonest indel (Fig. 7). Notably the magnitude of changes observed upon TSA treatment is comparable to those observed when inhibitors of specific DNA repair pathways are used (van Overbeek et al., 2016). These results show that the chromatin configuration of a given site contributes to defining its indel profile. Given the established role of chromatin in DNA repair (Kalousi and Soutoglou, 2016), and the involvement of multiple DNA repair pathways in mediating CRISPR-induced DNA editing (Maruyama et al., 2015; van Overbeek et al., 2016; Shou et al., 2018), altered recruitment of factors involved in different pathways may underlie the observed difference upon chromatin modulation.

In summary, our findings uncover general principles guiding CRISPR-mediated DNA editing in human cells and provide guidelines for a more effective and safer use of the technology, with important implications for clinical applications. They also reveal a striking influence of DNA sequence in dictating DSBs repair outcomes and lay the foundation for future mechanistic studies that can increase our understanding of end-joining processes in human cells.

## ACKNOWLEDGEMENTS

We thank the Crick Advanced Sequencing and Bioinformatics and Biostatistics facilities for preparing and sequencing NGS libraries and for help with data processing. We thank Her Majesty Queen Elizabeth II for starting the sequencing run containing our samples. We also thank Andrew J. Steele for helpful comments on the machine learning analysis. This work was supported by The Francis Crick Institute, which receives its core funding from Cancer Research UK (FC001110, FC001152), the UK Medical Research Council (FC001110, FC001152), and the Wellcome Trust (FC001110, FC001152), and by the CRUK Drug Discovery Award (C50796/A19448) provided by Cancer Research UK and Bayer Healthcare to PS. NML is a Winton Group Leader in recognition of the Winton Charitable Foundation’s support towards the establishment of the Francis Crick Institute. NML is additionally funded by a Wellcome Trust Joint Investigator Award (103760/Z/14/Z, the MRC eMedLab Medical Bioinformatics Infrastructure Award (MR/L016311/1) and core funding from the Okinawa Institute of Science & Technology Graduate University. This work was also supported by a Wellcome Trust PhD Training Fellowship for Clinicians Award (110292/Z/15/Z) to AMC.

## AUTHOR CONTRIBUTIONS

AMC performed most of the computational analysis and wrote the manuscript with PS. THB generated all reagents used in the study, performed the large-scale experiment and the ANN analysis. JM performed the chromatin modulation experiment. ARP and NML supervised the computational work and provided input on the manuscript. PS conceived the study, analysed the data, supervised the work and wrote the manuscript.

## MATERIALS AND METHODS

### sgRNAs pool generation

sgRNA pools were generated by combining equal volumes of saturated bacterial culture from the arrayed library described in (Henser-Brownhill et al., 2017), and extracting the resulting plasmid libraries. Six different pools were generated and independently transduced into HepG2 Cas9-expressing cells. This was necessary to reduce the library complexity and allow efficient detection of indels despite the high number of unedited sequences in the cell population – each sgRNA only infected a limited number of cells. We first tested three pools targeting 100 sites each (pools 100_1, 100_2 and 100_3). Once we confirmed efficient indel detection, we generated three sgRNA pools targeting 450 sites each (pools 450_5, 450_6, 450_7). 450 pools contained three distinct sgRNAs targeting the same 450 genes. 100 pools mainly contained sgRNAs present in the 450 pools with a few additional ones (Table S6). Although pools were transduced and processed independently, indel analysis was performed integrating data from the different pools. When assessing efficiency of indel formation, only data from 450 pools were used. This was done because indel counts for sgRNAs present in both 450 and 100 pools were artificially higher than those detected at sites targeted only with the 450 pools. When assessing editing precision, data from both 100 and 450 pools was combined, as frequencies of individual indels are not affected by differences in indel counts.

### Cell lines and viral transduction

HepG2 cells were cultured in Minimum Essential Media (MEM) with 10% FBS, and HEK-293T cells were cultured in Dulbecco’s Modified Eagle’s Medium (DMEM) with 10% FBS. All media was supplemented with 2mM L-glutamine, 100U/ml penicillin, and 100μg/ml streptomycin. HepG2 cells inducible for Cas9 expression were generated as previously described (Henser-Brownhill et al., 2017). Cas9 expression was induced with 1μg/mL doxycycline 1 day prior to infection with the sgRNAs and sustained until cells were harvested for genomic DNA extraction. Transduction of sgRNAs was performed using high titre virus, at an estimated MOI of at least 10, to increase the fraction of edited cells in the population for each sgRNA. To produce virus, 80% confluent HEK-293T cells were transfected with the sgRNA pools (pLenti_BSD_sgRNA plasmids), packaging plasmids (psPax2 and pMD2G) and pAdVantage at a ratio of 3:1 DNA to FugeneHD (Promega E2311). 24h after transfection viral particles were collected, filtered through a 0.45μm filter and used to infect Cas9-expressing HepG2 cells in the presence of 5μg/ml Polybrene. To increase infection efficiency, plated cells were spun for 2h at 2000rpm soon after the virus-containing supernatant was added. A second infection was carried out using viral particles collected 48h after transfection. Cells were not spun for the second infection. An sgRNA targeting the EZH2 gene was individually transduced in parallel, to allow determination of editing efficiency and assessment of possible phenotypic consequences of gene KO. Transduced cells were selected with 4μg/mL blasticidin, starting 24h after the first infection, and genomic DNA was extracted 5 days after infection (Qiagen 69506).

### Chromatin modulation

To investigate the effect of chromatin on CRISPR-mediated DNA editing, HepG2 cells were treated with the HDAC inhibitor Trichostatin A (Sigma), which induces histone hyperacetylation, and the EZH2 inhibitor GSK126 (Sigma), which globally reduces H3K27Me3 levels. Cells pre-treated with TSA (11nM or 100nM) or EZH2 (0.3 μM and 3 μM) for 5 days were infected with sgRNAs targeting the ACTL6A.5, ASF1B.7, BRD2.7, MBD3L1.6, MSH6.2 and SMARCD2.1 sites. Treatment was continued for an additional 5 days while indels were induced. Successful alteration of histone marks was confirmed by Western blot analysis.

### Protein detection

Western blot analysis and immunofluorescence microscopy were performed as previously described (Torres et al., 2016) using anti-pan Acetyl H3 (Abcam ab47915), anti-H3K27me3 (Millipore 07-449) and anti-EZH2 (CST 5246S) antibodies.

### Library preparation and NGS sequencing

#### sgRNA representation in pools

To assess the representation of individual sgRNAs in the plasmid library, amplicons containing the sgRNA sequences were generated as previously described (Henser-Brownhill et al., 2017). Briefly, PCR amplicons containing the P5 and P7 Illumina adaptors were generated using the high-fidelity Herculase II polymerase kit (Agilent 600675), and the resulting products extracted from an agarose gel (Qiagen 28706). Purified products were sequenced with either a HiSeq2500 or a MiSeq using custom sequencing and indexing primers (SeqP and IndexP, Table S7). Following sample demultiplexing, all sgRNA sequences were trimmed and aligned to the target sequences to assess sgRNA representation (normalized read count).

#### Large scale indel sequencing

To identify CRISPR-mediated editing at targeted regions, DNA libraries enriched for the targeted sites were generated using the SureSelect Target enrichment kit (Agilent) following the manufacturer’s instructions. Capture probes were designed to cover 2Kb regions centred on each target site. When multiple target sites were located in the same exon, the 2Kb region was centred on the exon middle point. Probe tiling parameters were: Tiling density: 1x; Masking: Least Stringent; Boosting: Maximize Performance. All samples were sequenced using Paired End 100bp runs on a HiSeq 4000 sequencer, multiplexing 2 samples per lane. Approximately 200 million reads were obtained for each sample. Analysis of sequenced regions confirmed good enrichment of the targeted regions (Fig. S2B).

#### Small scale indel sequencing

For validation experiments and experiments assessing the effect of chromatin modulation, indels induced at 6 selected sites were examined. In these experiments, a two-step PCR was performed on biological duplicates to generate a library of PCR amplicons. For the first PCR, 150ng of the corresponding gDNA were amplified for 20-22 cycles using the Herculase II polymerase kit yielding products of ~500bp (See Table S7 for primers). Next, PCR products were purified as per manufacturer’s instructions (Qiagen 28106) and 1μl of the resulting product was used as a template for the second nested PCR reaction in which primers containing barcodes and adapters for the sequencing reaction were added. Overall, a library of 60 individually barcoded amplicons of ~300bp was generated (See Table S7 for primers). Samples were purified in a 96-well format (Zymo Research D4018) and sequenced on a 300bp paired-end run on a MiSeq using standard Illumina sequencing primers (See Table S7 for primers). The long 300bp reads allowed assessment of both long and short indels.

### Sequenced read alignment

The quality of the sequenced reads was assured using FastQC (Andrews, 2010). We used BBMap (v. 36.59) (Bushnell, 2015) as it is a global aligner that is able to align longer indels. Alignment was carried out against the UCSC hg19/GRCh37 genome assembly.

### Indel identification

#### Large scale indel sequencing

In order to identify robustly the reads that contained indels we adopted a two-stage alignment strategy. In the first phase we aligned the reads to the genome disallowing any reads that contained indels. We discarded reads that aligned in a proper pair in this phase and took the remainder forward. In the second phase we aligned the remaining reads to the genome, this time setting a soft threshold permitting allowing indels up to 2000 bp. Duplicates were marked using Picard (v. 2.1.0). Reads that were marked as duplicates, or that had a mapping quality score of less than 38 were filtered using samtools (v. 1.2)(Li et al., 2009) and sambamba (v. 0.6.0)(Tarasov et al., 2015).

This two-phase approach was necessary to delineate, for a given target amplicon, between background arising from reads from cells uninfected with sgRNA and signal from reads from cells with successful transfections, on account of the pooling of sgRNAs. For a given amplicon, only a small proportion of the total number of cells would have been transfected with the sgRNA targeting the site contained within it. We know that aligned reads containing indels arise from appropriately transfected cells. However, our approach forces the aligner to choose an alignment with no indels over one with indels for the multiple possibilities for a given read. With this approach we can improve our confidence that the reads with indels are not background noise or alignment errors.

Indel identification was performed in R (v. 3.3.2 - 3.4.3) using custom scripts. The location and size of indels in reads were identified from the CIGAR string. Indels were only considered valid if they occurred within 5 nucleotides of the Cas9 cleavage site (defined as 6 nucleotides upstream of the end of the guide RNA including the PAM sequence). Any indels that could also be detected in the control HepG2 sample were removed as probable somatic mutations in this cancer cell line. We filtered out target sites that had indels identified in fewer than 10 reads (across all samples and replicates, where present) to ensure robust characterisation of these sites.

#### Small scale indel sequencing

Before alignment, paired end reads were merged using BBMerge (v. 36.59). After alignment, duplicates were marked using Picard (v. 2.1.0). Reads that were marked as duplicates, or that had a mapping quality score of less than 38 were filtered using samtools (v. 1.2) (Li et al., 2009) and sambamba (v. 0.6.0) (Tarasov et al., 2015). The R package CrispRVariants (Lindsay et al., 2016) was used to identify indels.

### Characterisation of target sites

Throughout, we used all detected indels from both 100 and 450 pools to characterise the targets, except when assessing for efficiency where indels from the 450 pools only were used to ensure an unbiased analysis of each target site as explained above.

#### Frameshifts and indel size

Indels were assessed for their frameshift potential by the divisibility of their size by 3. To identify patterns in the indel size profiles at target sites, we calculated the frequency of each size of indel (considered in bins of insertions greater than 1 nucleotide, insertions of 1, and deletions of 1, 2, 3, 4, 5, 6, 7, 8, 9, 10 and greater than 10). We performed unsupervised hierarchical clustering using the Ward D2 method to categorise groups of target sites based on their indel size profiles.

#### Precision

We also categorised target sites by calculated the frequency of each distinct indel at each target site. The most frequent indel was termed the commonest; ties were broken by prioritising insertions over deletions, and then by longest deletion. The precision of indel generation at a target site was defined based on the frequency of the commonest indel: imprecise ≤ 0.25, 0.25 < middle ≤ 0.5, precise > 0.5.

#### Sequence homology

The presence of microhomology of *n* nucleotides was assessed in the deletions. The 5’ *n* nucleotides of the deleted sequence were compared with the first *n* nucleotides downstream of the 3’ join. Likewise, the 3’ *n* nucleotides of the deleted sequence were compared with the last *n* nucleotides upstream of the 5’ join. If there was a match, this was considered as microhomology. For each deletion sequence, values of *n* ranging from 1 to 50 (or the length of the deletion, whichever was shortest) were used. The largest matching *n* was considered the size of the microhomology.

Insertion homology was assessed by extracting the inserted nucleotide from the read sequence using the CIGAR string. This was compared with the nucleotide in the −4 position of the protospacer to assess for matches. When assessing the commonest insertion, we only considered target sites that had 5 or more insertions. Where the inserted nucleotide either creates, or lies within a short repetitive stretch; e.g. “A” inserted adjacent to “A” creating “AA”, or “T” inserted adjacent to/within “TT” creating “TTT”; it is not possible to identify precisely which of these nucleotides is the inserted position. The aligner arbitrarily assigns the first position to the inserted nucleotide.

### Assessment of the effects induced by chromatin modulation

Mutation efficiency was assessed using the mutationEfficiency function from CrispRVariants (Lindsay et al., 2016), considering single nucleotide variants as non-variants. To compare the counts of indels across the different conditions, in order to assess the contribution of each indel to the changes in efficiency, the raw counts for each indel in each condition were normalised to the library size for that condition. Indels that constituted less than 1% of the library size in any condition were filtered out.

To assess the effects of chromatin modulation on the indel profile of target, over and above the effects on efficiency, we performed a different normalisation on the raw counts. We divided by a size factor (the total number of indels detected in a condition). In this way, we could compare the relative contribution of each indel to the overall indel profile across the different conditions. After normalisation, only the most frequent 10 indels in the untreated condition were used.

### Analysis of chromatin environment

DNase-seq and H3K9ac and H3K27ac ChIP-seq fold-enrichment data for in HepG2 cells were obtained pre-processed from the Roadmap Epigenomics consortium (Roadmap Epigenomics Consortium et al., 2015). We calculated the mean fold-enrichment signal in a 500 bp window centred on the cleavage site of the six validation targets.

### Precision prediction and feature influence detection

To predict precision, we designed an artificial neural network (ANN) that uses the raw sgRNA sequences as input: 20 individual nucleotides, plus the PAM sequence (as a rudimentary internal control). All variable nucleotides were encoded using one hot encoding. The input layer of the network therefore has 86 nodes, with each position represented by 4 binary inputs (except the constant ’GG’ in the PAM sequence). These are followed by a single hidden layer containing 512 neurons using rectified linear unit (ReLU) activation functions, connected to a single output node, followed by a softplus activation function. Our loss function was mean square error (L2 norm loss). The model parameters were initialised using Xavier initialisation. In summary, the weights were initially filled with random numbers [−c, c] where:

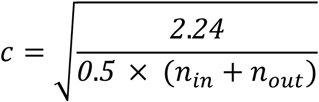

Here, *n_in_*, *n_in_* is the number of neurons preceding weights and *n_out_* is the number of neurons proceeding weights. The ANN was trained for 1500 epochs using stochastic gradient descent with Nesterov momentum set to 0.9, a learning rate of 0.001, and a batch size of 100. To ensure consistency and to reduce bias introduced by particular sets of sgRNAs, we performed bootstrapping with replacement, training the ANN with our 646 sgRNAs randomly divided into training (n = 517 (80%)) and validation (n = 129 (20%)) sets 10 times. To identify key sequence positions with the greatest influence on predicting precision, we conducted a permutation “nucleotide” importance analysis by systematically neutralising each nucleotide at the respective position by setting it to zero in every validation sgRNA, then recording the reduction in correlation coefficients (linear R^2^) from predictions made for the unaltered sequences. We also recorded the difference in predictive statistical significance (Wald test p-values). We performed this procedure for all 10 trained ANNs and report the average percentage reduction in R^2^ for neutralised positions. We considered an increase in average cross-model Wald test p value > 0.05 as having abolished the predictive significance of the model. The ANN was built, trained, and deployed using *Apache MXNET* (python 3 API) v. 1.2.0.

### Statistical and computational analysis

Non-parametric statistical tests were used and p-values were adjusted for multiple testing where necessary; each specific test is indicated in the text. Unless otherwise stated, all downstream bioinformatic analysis was performed in R (v. 3.3.2 - 3.4.3). Custom scripts are available at https://github.com/luslab/crispr-indels.

### Data availability

Sequencing data are available on EBI ArrayExpress at accessions E-MTAB-7091 and E-MTAB-7095.

## SUPPLEMENTARY FIGURE LEGENDS

**Figure S1.**
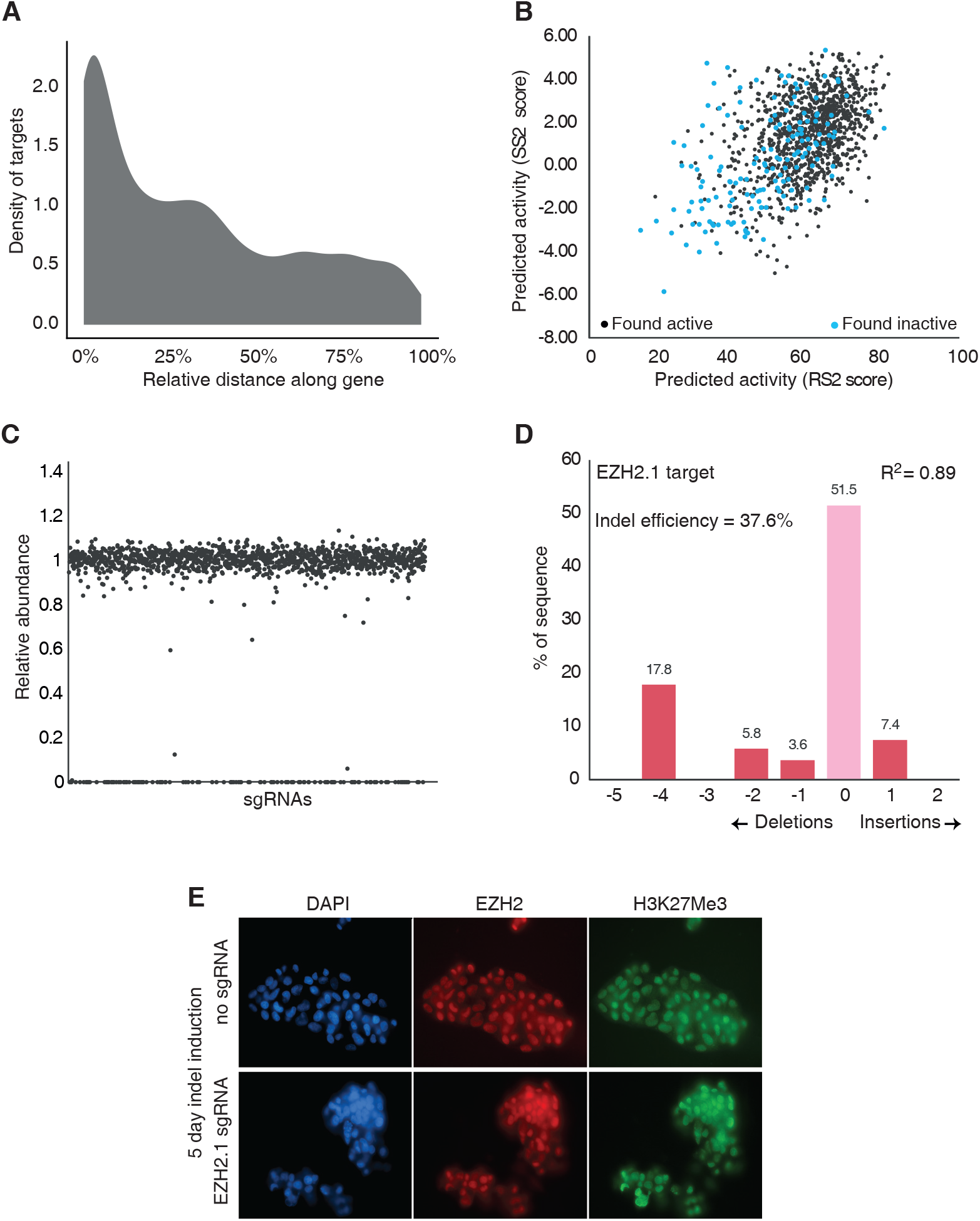
Characteristics of the sgRNA pools. **A**. The distribution of cleavage site locations of all target sites along the gene body is shown. **B**. Correlation between the indicated scores of predicted sgRNA activity for the used guides obtained from two distinct algorithms (Chari et al., 2017; Doench et al., 2016). sgRNAs that induced detectable indels are shown in black, while inactive sgRNAs are in blue. Note that several sgRNAs with high predicted activity were found to be inactive. **C**. Relative abundance of each sgRNA in the 450 pools. Values represent sequencing reads normalized to the median value of the three pools combined. With the exception of a few undetectable sgRNAs, all guides are homogeneously represented in the pools. **D**. TIDE analysis (Brinkman et al., 2014) examining the editing efficiency at the EZH2.1 target, used as a reporter of indel formation efficiency in the large-scale experiment. EZH2.1 sgRNA was individually transduced in Cas9-expressing cells using the same conditions used for the pooled sgRNAs. Output from the TIDE website https://tide.nki.nl/, focusing on the most reliable indels (p <0.001), is shown, indicating good indel efficiency 5 days after transduction. **E**. Immunofluorescence microscopy of the indicated samples using anti-EZH2 (red) and anti-H3K27Me3 (green) antibodies, showing no detectable reduction in EZH2 or histone modification levels 5 days after indel induction. Nuclei are counterstained with DAPI (blue).

**Figure S2.**
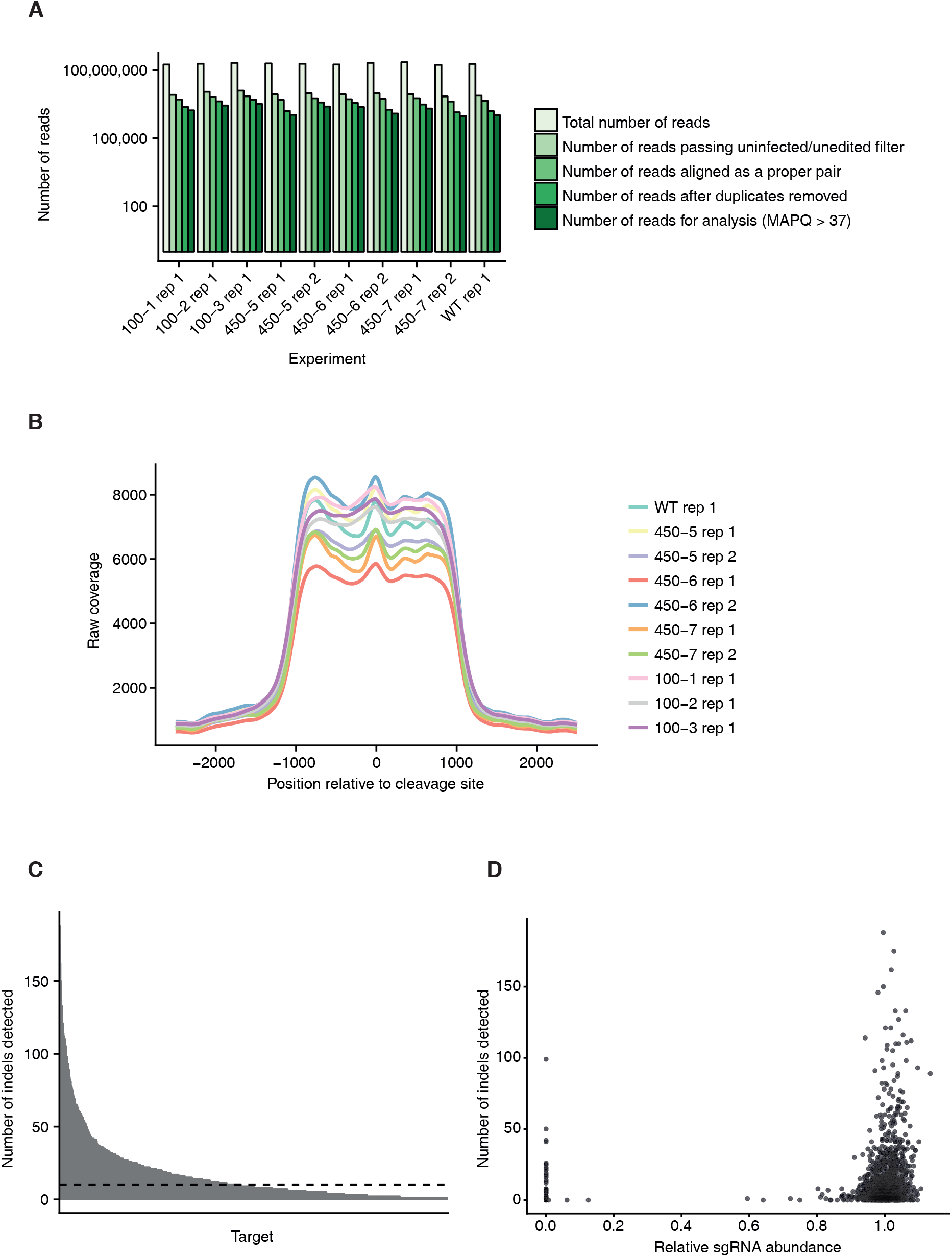
Indel detection metrics. **A**. Alignment metrics for the large-scale experiments. For each experiment, the number of reads at each stage of processing is shown. The filtering strategy used to detect indels robustly is described in the Methods section. **B**. The raw coverage of the phase 1 reads (those without indels detected) for each experiment is shown over the region around the cleavage site that was selectively isolated by target capture. **C**. The number of total indels detected at each target site when using the 450 sgRNA pools is shown. The dashed line is at 10, reflecting the threshold that was set for the downstream analysis. **D**. The relationship between the number of total indels detected at each target site and the abundance of the associated sgRNA in the pools is shown. Note that some sgRNAs that were undetectable in the pools by next-generation sequencing induced indels at their target sites. Presence of the undetected, indel-inducing sgRNAs was confirmed by Sanger sequencing of the individual guides in the original arrayed library.

**Figure S3.**
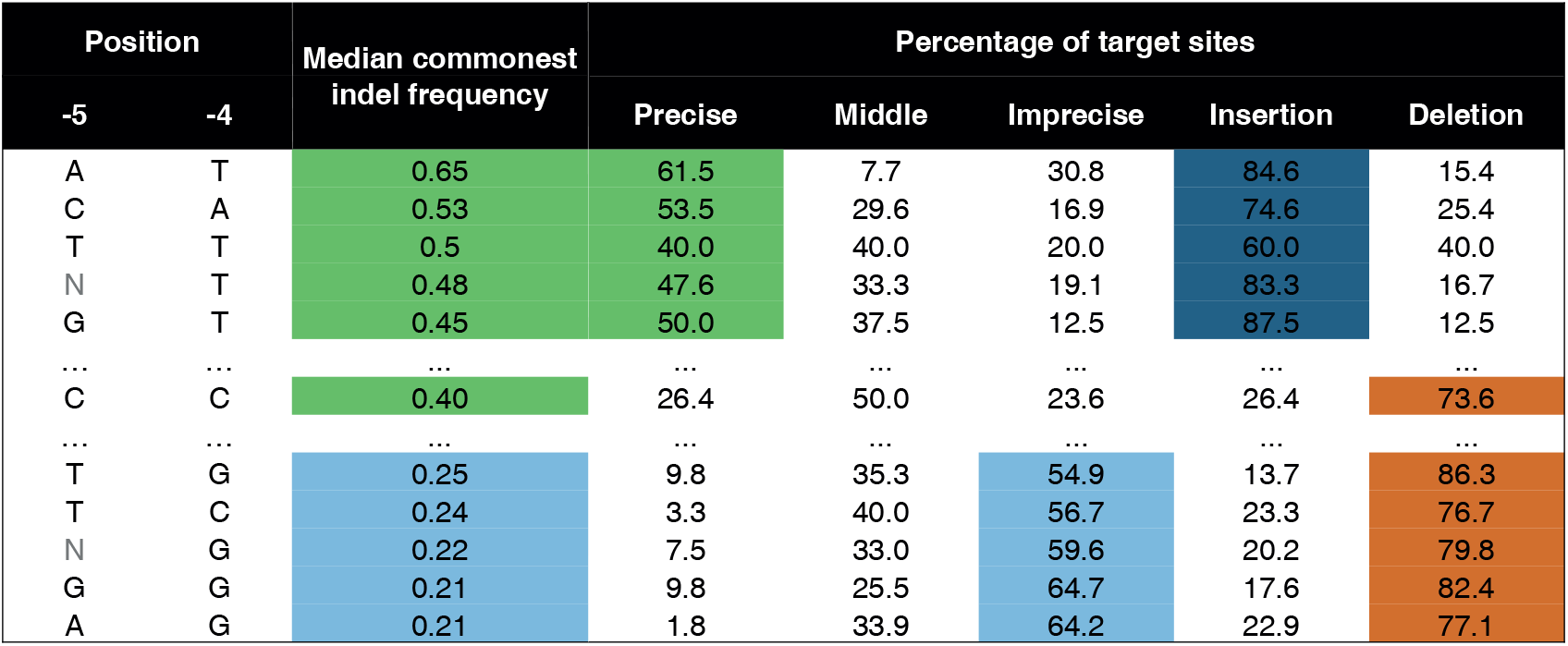
Predictive nucleotides within the protospacer. Associated with Fig. 5. Precision rates and insertion or deletion preferences for nucleotide combinations either considering the −5 and −4 positions together, or the −4 position alone. Single nucleotides or di-nucleotides that best predict precise targets with a preference for insertions (green and dark blue), precise targets with a preference for deletions (green and orange) or imprecise targets with a preference for deletions (light blue and orange) are shown. Rows with dots indicate other nucleotide combinations that showed a weaker correlation with these features. Nucleotide combinations are ranked based on the median commonest indel frequency.

**Figure S4.**
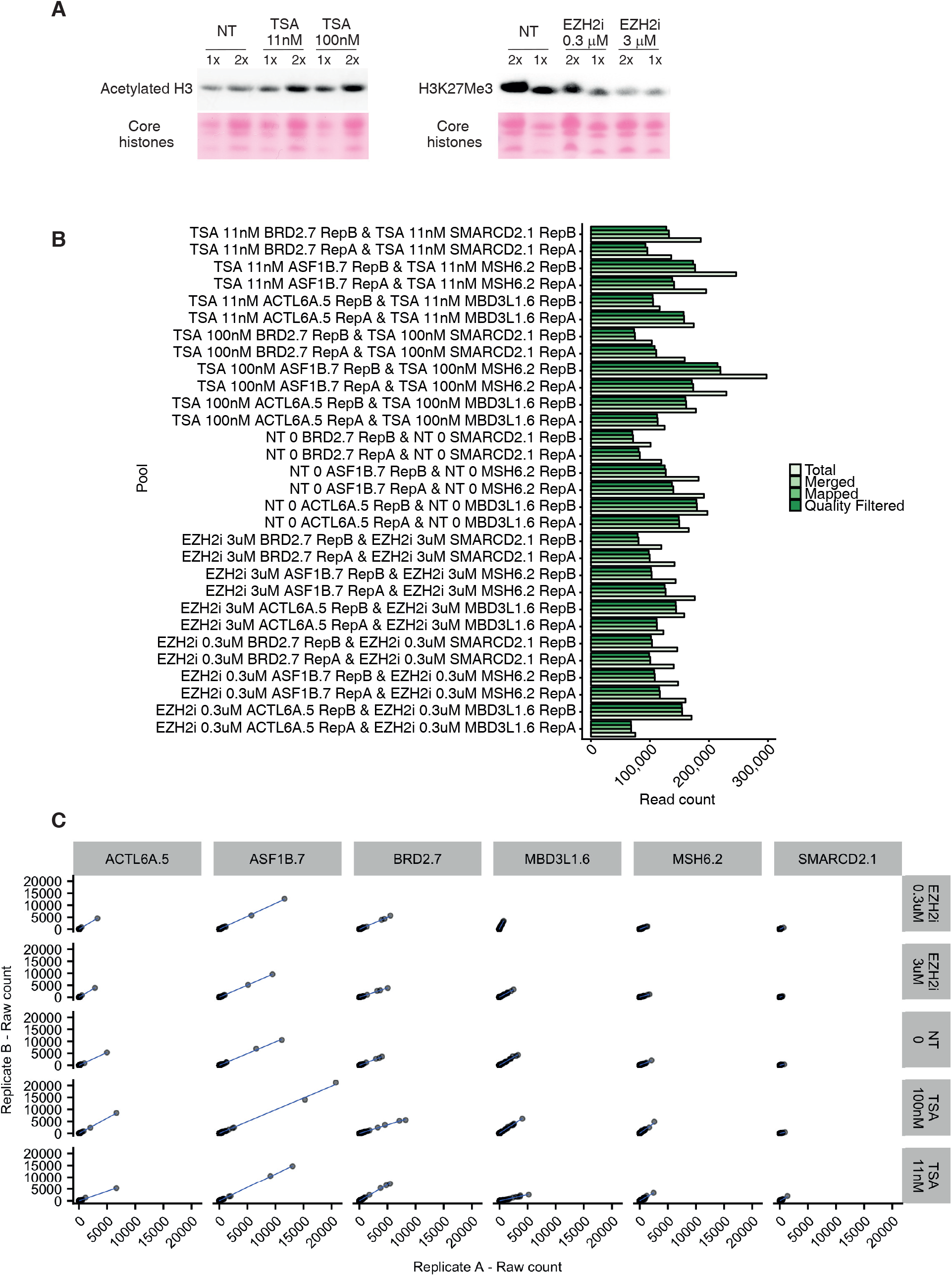
Indel detection upon chromatin modulation. **A**. Western blot analysis of transduced HepG2 cells untreated (NT) or treated with the indicated compounds. A dose-dependent increase in acetylated H3 is observed in response to TSA (left), while the EZH2i induces a dose-dependent reduction in H3K27Me3 (right). Ponceau staining of the core histones is shown as loading control. Two different amounts of protein lysate (1x and 2x) are loaded for each condition to allow a more quantitative assessment of the differences. **B**. The alignment metrics for the experiments testing the effect of chromatin on indel profiles. For each experiment, the number of reads at each stage of processing is shown. **C**. Correlation between raw read counts (prior to normalisation) for each detected indel between both biological replicates is shown for each condition.

**Figure S5.**
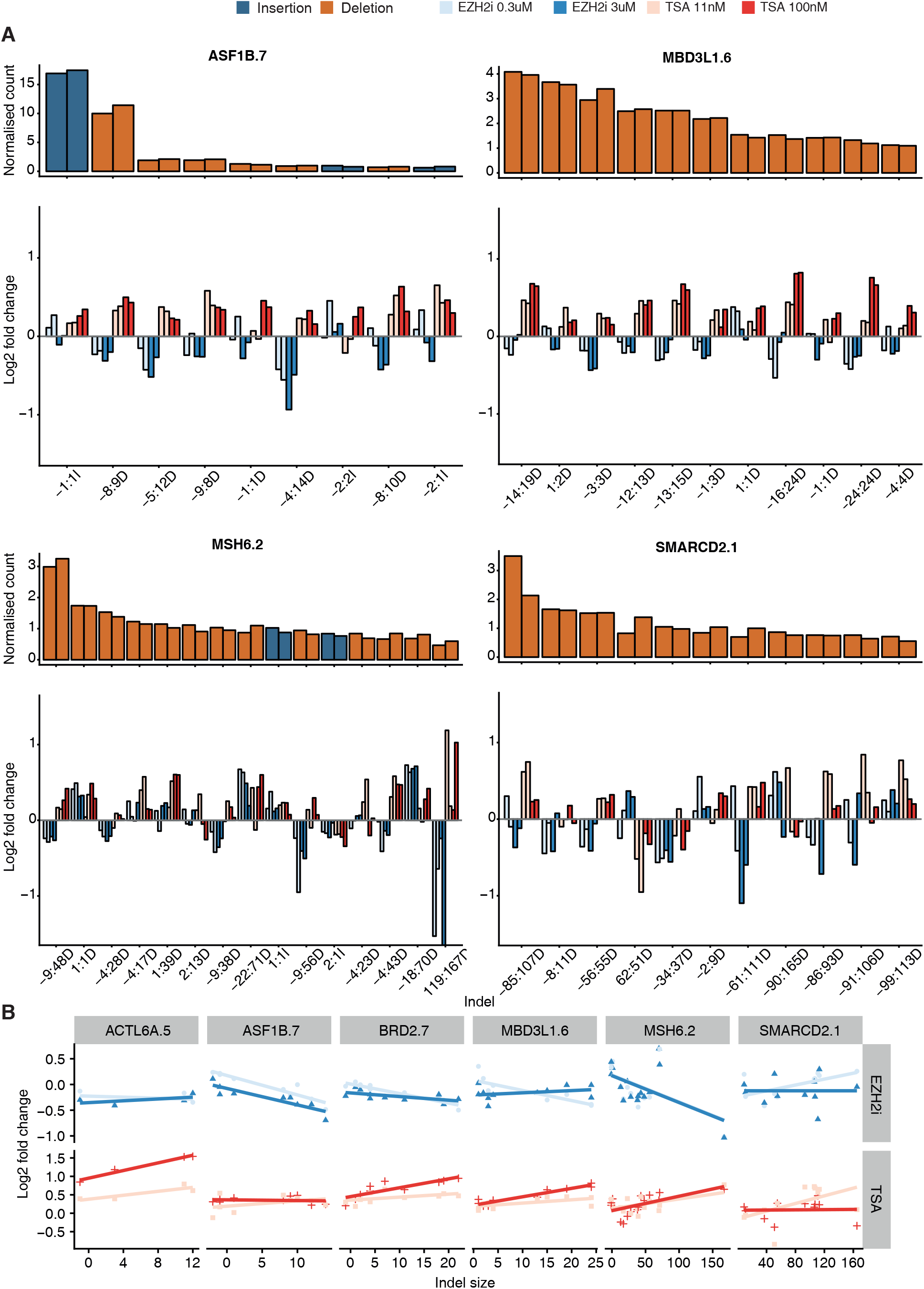
Chromatin modulation affects RGN activity. **A**. As for Figure 6D, the remaining 4 target sites show that chromatin modulation affects both insertions and deletions. Above is shown the count of each indel, normalised by the effective library size at each site for each replicate. Only indels with a normalised count of at least 1 in any condition are included. **B**. The log2-fold change for each size of indel is shown for each target site and for each condition.

**Figure S6.**
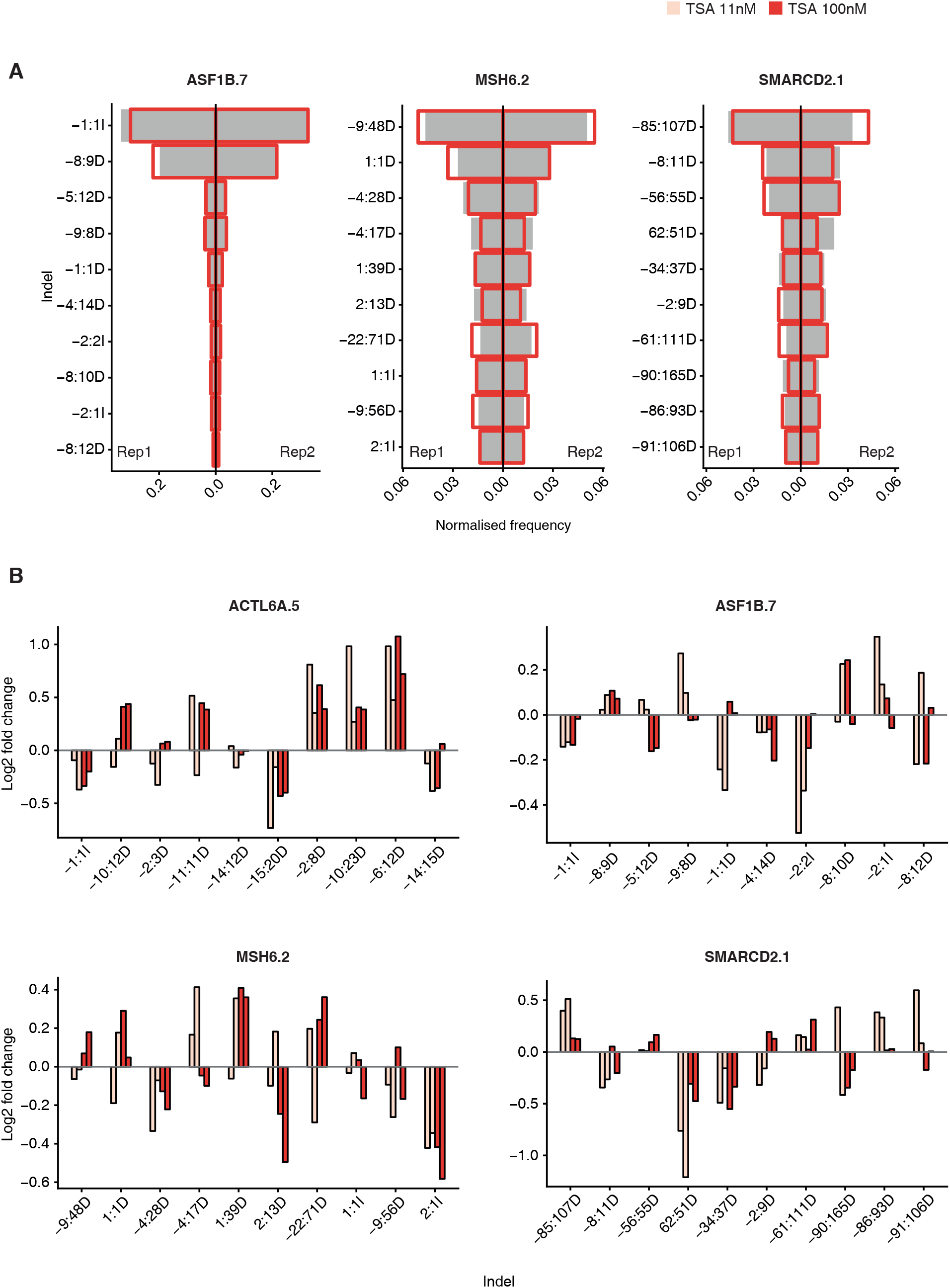
Chromatin modulation affects indel profiles. **A**. As for Fig. 7A, the remaining 3 targets show that chromatin modulation has a differential effect on distinct indels. The count for each indel, normalised by the total number of indels detected at that target site in that condition is shown for each replicate. The frequency of the indicated indels in the untreated condition (grey bars) and in the TSA 100nM condition (red outline) is shown. The 10 commonest indels for each site are shown. **B**. As for Fig. 7B, the log2 fold change in frequency for the indicated indels is shown for the remaining 4 target sites. The 10 commonest indels across both replicates are shown.

## REFERENCES

Andrews, S. (2010). FastQC: a quality control tool for high throughput sequence data. https://www.bioinformatics.babraham.ac.uk/projects/fastqc/

Bae, S., Kweon, J., Kim, H.S., and Kim, J.-S. (2014). Microhomology-based choice of Cas9 nuclease target sites. Nat. Methods 11, 705–706.

Brinkman, E.K., Chen, T., Amendola, M., and van Steensel, B. (2014). Easy quantitative assessment of genome editing by sequence trace decomposition. Nucleic Acids Res. 42, e168.

Brinkman, E.K., Chen, T., de Haas, M., Holland, H.A., Akhtar, W., and van Steensel, B. (2018). Kinetics and Fidelity of the Repair of Cas9-Induced Double-Strand DNA Breaks. Mol. Cell 70, 801–813.e6.

Bushnell, B. (2015). BBMap. https://sourceforge.net/projects/bbmap/

Deriano, L., and Roth, D.B. (2013). Modernizing the nonhomologous end-joining repertoire: alternative and classical NHEJ share the stage. Annu. Rev. Genet. 47, 433–455.

Doench, J.G., Fusi, N., Sullender, M., Hegde, M., Vaimberg, E.W., Donovan, K.F., Smith, I., Tothova, Z., Wilen, C., Orchard, R., et al. (2016). Optimized sgRNA design to maximize activity and minimize off-target effects of CRISPR-Cas9. Nat. Biotechnol. 34, 184–191.

Dudley, D.D., Chaudhuri, J., Bassing, C.H., and Alt, F.W. (2005). Mechanism and control of V(D)J recombination versus class switch recombination: similarities and differences. Adv. Immunol. 86, 43–112.

Geisinger, J.M., Turan, S., Hernandez, S., Spector, L.P., and Calos, M.P. (2016). In vivo blunt-end cloning through CRISPR/Cas9-facilitated non-homologous end-joining. Nucleic Acids Res. 44, e76.

van Heemst, D., Brugmans, L., Verkaik, N.S., and van Gent, D.C. (2004). End-joining of blunt DNA double-strand breaks in mammalian fibroblasts is precise and requires DNA-PK and XRCC4. DNA Repair 3, 43–50.

Henser-Brownhill, T., Monserrat, J., and Scaffidi, P. (2017). Generation of an arrayed CRISPR-Cas9 library targeting epigenetic regulators: from high-content screens to in vivo assays. Epigenetics 12, 1065–1075.

Horlbeck, M.A., Witkowsky, L.B., Guglielmi, B., Replogle, J.M., Gilbert, L.A., Villalta, J.E., Torigoe, S.E., Tjian, R., and Weissman, J.S. (2016). Nucleosomes impede Cas9 access to DNA in vivo and in vitro. Elife 5.

Hsu, P.D., Lander, E.S., and Zhang, F. (2014). Development and applications of CRISPR-Cas9 for genome engineering. Cell 157, 1262–1278.

Isaac, R.S., Jiang, F., Doudna, J.A., Lim, W.A., Narlikar, G.J., and Almeida, R. (2016). Nucleosome breathing and remodeling constrain CRISPR-Cas9 function. Elife 5, e13450

Jensen, K.T., FlØe, L., Petersen, T.S., Huang, J., Xu, F., Bolund, L., Luo, Y., and Lin, L. (2017). Chromatin accessibility and guide sequence secondary structure affect CRISPR-Cas9 gene editing efficiency. FEBS Lett. 591, 1892–1901.

Jinek, M., Chylinski, K., Fonfara, I., Hauer, M., Doudna, J.A., and Charpentier, E. (2012). A programmable dual-RNA-guided DNA endonuclease in adaptive bacterial immunity. Science 337, 816–821.

Kalousi, A., and Soutoglou, E. (2016). Nuclear compartmentalization of DNA repair. Curr. Opin. Genet. Dev. 37, 148–157.

Kosicki, M., Tomberg, K., and Bradley, A. (2018). Repair of double-strand breaks induced by CRISPR-Cas9 leads to large deletions and complex rearrangements. Nat. Biotechnol. dx.doi.org/10.1038/nbt.4192

Lemos, B.R., Kaplan, A.C., Bae, J.E., Ferrazzoli, A.E., Kuo, J., Anand, R.P., Waterman, D.P., and Haber, J.E. (2018). CRISPR/Cas9 cleavages in budding yeast reveal templated insertions and strand-specific insertion/deletion profiles. Proc. Natl. Acad. Sci. U. S. A. 115, E2040–E2047.

Li, H., Handsaker, B., Wysoker, A., Fennell, T., Ruan, J., Homer, N., Marth, G., Abecasis, G., Durbin, R., and 1000 Genome Project Data Processing Subgroup (2009). The Sequence Alignment/Map format and SAMtools. Bioinformatics 25, 2078–2079.

Lindsay, H., Burger, A., Biyong, B., Felker, A., Hess, C., Zaugg, J., Chiavacci, E., Anders, C., Jinek, M., Mosimann, C., et al. (2016). CrispRVariants charts the mutation spectrum of genome engineering experiments. Nat. Biotechnol. 34, 701–702.

Maruyama, T., Dougan, S.K., Truttmann, M.C., Bilate, A.M., Ingram, J.R., and Ploegh, H.L. (2015). Increasing the efficiency of precise genome editing with CRISPR-Cas9 by inhibition of nonhomologous end joining. Nat. Biotechnol. 33, 538–542.

van Overbeek, M., Capurso, D., Carter, M.M., Thompson, M.S., Frias, E., Russ, C., Reece-Hoyes, J.S., Nye, C., Gradia, S., Vidal, B., et al. (2016). DNA Repair Profiling Reveals Nonrandom Outcomes at Cas9-Mediated Breaks. Mol. Cell 63, 633–646.

Roadmap Epigenomics Consortium, Kundaje, A., Meuleman, W., Ernst, J., Bilenky, M., Yen, A., Heravi-Moussavi, A., Kheradpour, P., Zhang, Z., Wang, J., et al. (2015). Integrative analysis of 111 reference human epigenomes. Nature 518, 317–330.

Shou, J., Li, J., Liu, Y., and Wu, Q. (2018). Precise and Predictable CRISPR Chromosomal Rearrangements Reveal Principles of Cas9-Mediated Nucleotide Insertion. Mol. Cell dx.doi.org/10.1016/j.molcel.2018.06.021.

Taheri-Ghahfarokhi, A., Taylor, B.J., Nitsch, R., Lundin, A., Cavallo, A.-L., Madeyski-Bengtson, K., Karlsson, F., Clausen, M., Hicks, R., Mayr, L.M., et al. (2018). Decoding non-random mutational signatures at Cas9 targeted sites. Nucleic Acids Res. dx.doi.org/10.1093/nar/gky653

Tarasov, A., Vilella, A.J., Cuppen, E., Nijman, I.J., and Prins, P. (2015). Sambamba: fast processing of NGS alignment formats. Bioinformatics 31, 2032–2034.

Torres, C.M., Biran, A., Burney, M.J., Patel, H., Henser-Brownhill, T., Cohen, A.-H.S., Li, Y., Ben-Hamo, R., Nye, E., Spencer-Dene, B., et al. (2016). The linker histone H1.0 generates epigenetic and functional intratumor heterogeneity. Science 353, aaf1644.

Tsai, S.Q., Zheng, Z., Nguyen, N.T., Liebers, M., Topkar, V.V., Thapar, V., Wyvekens, N., Khayter, C., Iafrate, A.J., Le, L.P., et al. (2015). GUIDE-seq enables genome-wide profiling of off-target cleavage by CRISPR-Cas nucleases. Nat. Biotechnol. 33, 187–197.

Uusi-Mäkelä, M.I.E., Barker, H.R., Bauerlein, C.A., Hakkinen, T., Nykter, M., and Ramet, M. (2018). Chromatin accessibility is associated with CRISPR-Cas9 efficiency in the zebrafish (Danio rerio). PLoS One 13, e0196238.

Wang, T., Wei, J.J., Sabatini, D.M., and Lander, E.S. (2014). Genetic screens in human cells using the CRISPR-Cas9 system. Science 343, 80–84.

## References

Chari, R., Yeo, N.C., Chavez, A., and Church, G.M. (2017). sgRNA Scorer 2.0: A Species-Independent Model To Predict CRISPR/Cas9 Activity. ACS Synth. Biol. 6, 902–904.

